# A subset of Memory B-derived antibody repertoire from 3-dose vaccinees is ultrapotent against diverse and highly transmissible SARS-CoV-2 variants, including Omicron

**DOI:** 10.1101/2021.12.24.474084

**Authors:** Kang Wang, Zijing Jia, Linlin Bao, Lei Wang, Lei Cao, Hang Chi, Yaling Hu, Qianqian Li, Yinan Jiang, Qianhui Zhu, Yongqiang Deng, Pan Liu, Nan Wang, Lin Wang, Min Liu, Yurong Li, Boling Zhu, Kaiyue Fan, Wangjun Fu, Peng Yang, Xinran Pei, Zhen Cui, Lili Qin, Pingju Ge, Jiajing Wu, Shuo Liu, Yiding Chen, Weijin Huang, Cheng-Feng Qin, Youchun Wang, Chuan Qin, Xiangxi Wang

## Abstract

Omicron, the most heavily mutated SARS-CoV-2 variant so far, is highly resistant to neutralizing antibodies, raising unprecedented concerns about the effectiveness of antibody therapies and vaccines. We examined whether sera from individuals who received two or three doses of inactivated vaccine, could neutralize authentic Omicron. The seroconversion rates of neutralizing antibodies were 3.3% (2/60) and 95% (57/60) for 2- and 3-dose vaccinees, respectively. For three-dose recipients, the geometric mean neutralization antibody titer (GMT) of Omicron was 15, 16.5-fold lower than that of the ancestral virus (254). We isolated 323 human monoclonal antibodies derived from memory B cells in 3-dose vaccinees, half of which recognize the receptor binding domain (RBD) and show that a subset of them (24/163) neutralize all SARS-CoV-2 variants of concern (VOCs), including Omicron, potently. Therapeutic treatments with representative broadly neutralizing mAbs individually or antibody cocktails were highly protective against SARS-CoV-2 Beta infection in mice. Atomic structures of the Omicron S in complex with three types of all five VOC-reactive antibodies defined the binding and neutralizing determinants and revealed a key antibody escape site, G446S, that confers greater resistance to one major class of antibodies bound at the right shoulder of RBD through altering local conformation at the binding interface. Our results rationalize the use of 3-dose immunization regimens and suggest that the fundamental epitopes revealed by these broadly ultrapotent antibodies are a rational target for a universal sarbecovirus vaccine.

**One sentence summary:** A sub-set of antibodies derived from memory B cells of volunteers vaccinated with 3 doses of an inactivated SARS-CoV-2 vaccine work individually as well as synergistically to keep variants, including Omicron, at bay.

## Main Text

The ongoing evolution and emergence of severe acute respiratory syndrome coronavirus 2 (SARS-CoV-2) variants raise concerns about the effectiveness of monoclonal antibodies (mAbs) therapies and vaccines ^1–3^, posing challenges for global pandemic control. These variants were characterized as Variant of Interest, VOI or Variant of Concern, VOC by the World Health Organization (WHO). The more recently identified Omicron variant (B.1.1.529), designated as a new VOC, has led to an unprecedented surge in COVID-19 cases in South Africa and is now spreading across the world ^4^. Remarkably, Omicron is the most heavily mutated variant to emerge so far with over thirty mutations in spike (S) protein, fifteen of which occur in the receptor binding domain (RBD). In addition, there are three small deletions and one 3-residue insertion in the N-terminal domain (NTD) of S1 subunit (Fig. 1a). The pattern of some of these alterations, similar to the those noted in previous VOCs, such as Δ69-70 in Alpha, N501Y in Alpha, Beta and Gamma, P681H in Alpha and Delta, are presumably associated with enhanced transmissibility, while many substitutions, including G142D/Δ143-145, ins214EPE, K417N, T478K, E484A, Q493K and N501Y, are closely related with resistance to neutralizing antibodies and vaccine induced humoral immunity ^1,3,5–9^ (Figs. 1a and 1b).

**Fig. 1.**
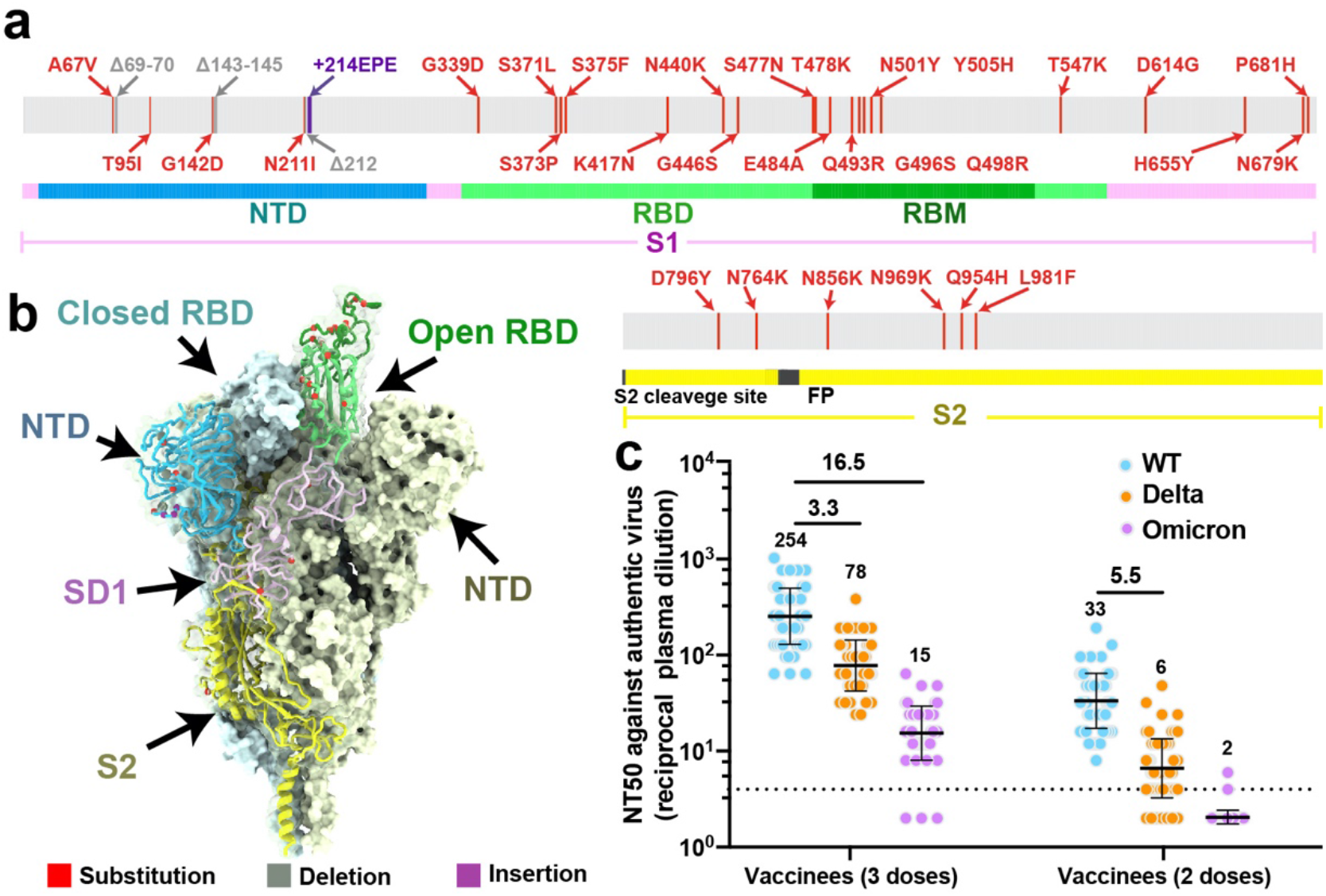
Evolution and neutralization characteristics of Omicron variant. **a**,, A linear representation of Omicron spike with mutations marked on. The replacements are marked in red; deletions are in grey and insertions are in purple. **b**, Distribution of mutations of Omicron on the cryo-EM structure of pre-fusion spike trimer. The mutations listed in **a** are indicated in the ‘up’ protomer shown as cartoon with mutated residues highlighted as spheres and colored as in **a**. The RBD, NTD, SD1 and S2 of this subunit are marked with arrow and colored in green, blue, magenta and yellow, respectively; the other two protomers in ‘down’ state are shown as surface in pale cyan and pale yellow, respectively. **c**, Graph shows the neutralizing antibody response against wild-type and Omicron SARS-CoV-2 authentic virus for sera from healthy vaccinees who received two doses (n=60) or three doses (n=60) of Coronavac. Neutralizing antibody titer fold decline for Delta or Omicron over wild-type for each group of sera is shown in each of the plots.

Although COVID-19 vaccines continued to be effective against severe diseases and deaths, including those caused by the circulating Delta variant, waning immunity and massive breakthrough infections caused by viral diversification warrant the need for a third dose or new vaccines. To combat the current resurgence of the epidemic, the U.S. Food and Drug Administration has authorized use of a 3^rd^ booster dose for all adults after completion of primary vaccination with approved COVID-19 vaccine ^10^. This step seems essential because preliminary studies have indicated that three doses of Pfizer-BioNtech mRNA vaccine neutralize the Omicron variant with an approximate 40-fold decline, while two doses are less effective ^11,12^. However, these preliminary data on the neutralization sensitivity of Omicron require further independent confirmation. The clinical impact of natural and vaccine-induced immunity with regards to protection from infection and severe disease needs urgent investigation.

## Authentic virus neutralization of the Omicron variant by vaccine sera

The CoronaVac, a β-propiolactone-inactivated vaccine against COVID-19, has been approved for emergency use, and recommended for a booster dose (third) of inactivated vaccine in older persons by WHO ^13,14^. Serum specimens from two groups of 2-dose (n=60, at month 0, 1) or 3-dose (n=60, at months 0, 1, 7) CoronaVac vaccinee volunteers were collected for evaluating neutralization titers against the Omicron and Delta variants using a live SARS-CoV-2. None of the volunteers recruited for vaccination was infected by SARS-CoV-2 prior to the study. Blood samples from vaccinees collected 4 weeks after the last vaccination were used in this study, to compare NAb titers against circulating SARS-CoV-2 variants. An early passage of isolated (CHK06 strain) and sequence confirmed live Omicron virus was used for neutralization assay in this study. Among three doses of CoronaVac recipients, the geometric mean half-maximal neutralizing titers (GMT NT_50_) against live wild type (WT) virus, Delta and Omicron variants were 254, 78 and 15, respectively. Compared with WT, neutralizing titers against Delta and Omicron were, on average, 3.3-fold and 16.5-fold reduced, respectively (Fig. 1c). Only 3 of 60 samples had a NT_50_ titer of < 8 against the Omicron with a seroconversion rate of 95% for neutralizing antibodies (Fig. 1c). However, it’s more concerning about effectiveness for two-dose regime against Omicron infection. Among two doses of CoronaVac recipients, NT_50_ titer against Delta was 6.3 with a 5-fold reduction when compared to WT, but none of the serum specimens had an NT_50_ titer of >8 against Omicron (Fig. 1c). Compared to 2-dose vaccinees, sera of the 3-dose vaccinees displayed lower reduction in neutralization titers against Delta, which is consistent with previous observations that 3-dose administration of inactivated vaccine leads to enhanced neutralizing breadth to SARS-CoV-2 variants ^5^.

## Three doses of vaccine-elicited monoclonal antibodies

We previously sorted immunoglobulin (IgG+) memory B cells from peripheral blood mononuclear cells (PBMCs) of four 3-dose CoronaVac vaccinees using prefusion SARS-CoV-2 S as a bait ^5,15^. In total, we sorted 1800 SARS-CoV-2 S-specific memory B cells, obtained 422 paired heavy- and light-chain antibody sequences, and selected 323 antibodies for expression. Characterization by ELISA showed that 163, 100 and 51 recognized the RBD, NTD and S2, respectively and 9 failed to bind S (Fig. 2a). Biolayer interferometry affinities (BLI) measurements showed that nearly all RBD-directed antibodies bound to WT SARS-CoV-2 at sub-nM levels and 127 of them showed neutralization activities against both authentic and pseudotyped WT SARS-CoV-2 were selected for further investigation. Of these antibodies, over 93% of these antibodies exhibited broad binding activities to most VOCs and VOIs. Notably, 85% of these antibodies cross-reacted with the Omicron RBD. Contrarily, ~80% of NTD antibodies lost their associations with Omicron. Additionally, NTD antibodies also showed relatively poor cross-reactivity to other four VOCs due to the greater diversity of the NTD (Fig. 1, a, b).

**Fig. 2.**
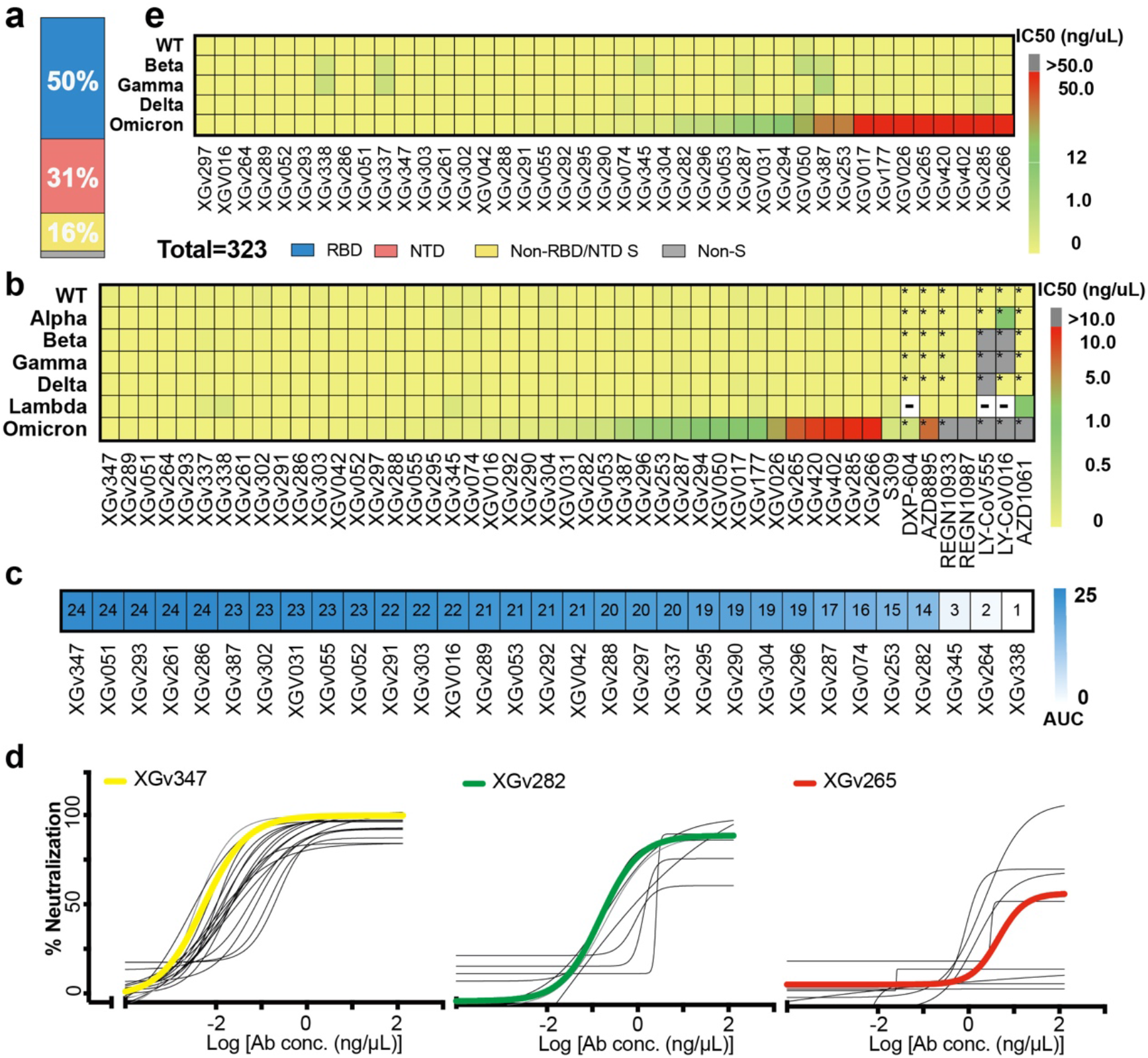
Characteristics of a subset of broadly neutralizing antibodies from recipients of a booster immunization. **a,** Vertical slices chart shows the gross binding epitope distribution of mAbs isolated from the individuals who received three doses of inactivated vaccines. Total number of antibodies and the percentage of antibodies that recognize RBD (blue), NTD (red) and S2 domain (yellow) are indicated. **b**, Heatmap representation of 41 selected representative mAbs and another 9 mAbs approved by FDA or in clinical trials against pseudotyped viruses with wild-type or variant SARS-CoV-2 S. The color bar on the right represents the ranges of IC_50_ values for the indicated mAbs against pseudotyped viruses in **c** (yellow: 0.002-0.020 ng/μL; green: 0.020-1.000 ng/μL; red: 1.000-10.000 ng/μL). Data marked with ‘*’ represents the data referred from the available publication in which the whole experiment system and condition of pseudovirus neutralization assay remains the samewith this study ^18^; Data marked with means no related datasets here. **c**, Heatmap with values shown in the form of AUC represents the competition ability between the selected mAbs and hACE2. Color gradient ranging from white (1) to blue (24) is shown on the right represents the competition ability from the weakest to the strongest. **d**, Neutralization curves for the selected 41 antibodies on pseudotyped viruses with the S protein of Omicron variant of concern. Data shown here are three groups of antibodies: 1) ultrapotent antibodies against all five VOCs, 2) highly potent antibodies against other four VOCs, but with median neutralizing activities against Omicron, 3) highly potent antibodies against other four VOCs, but with weak neutralizing activities against Omicron. XGv347, XGv282 and XGv265, selected as a representative of each group are highlighted by bold curve in yellow, green, and red, respectively, in correspondence with the color range in **e**. Heatmap representation of the same 41 mAbs as in **b** against wild-type and variant SARS-CoV-2 authentic viruses with color gradient shown on the right.

## A subset of antibodies with broad neutralization against circulating variants

Results of the pseudovirus neutralization assays performed by carrying the S of WT or other VOCs ^16,17^ identified XX RBD targeting antibodies that were especially potent with their half-maximal inhibitory concentration (IC_50_) ranging from 0.002 to 0.800 ng/μl against WT as well as all VOCs (Fig. 2). Among these, 28 antibodies executed their neutralization via directly blocking the interactions between the RBD and its receptor hACE2, while 3 antibodies employ other mechanisms to neutralize viral infection (Fig. 2d, Extended Data Fig. 1). Especially, a subset of RBD antibodies (13 and 24) neutralized Omicron with IC_50_ < 0.02 and 0.1 ng/μl, respectively. These neutralizations are as potent as those exhibited by best-in-class antibodies against WT (Fig. 2b and 2d). We obtained IC_50_ values of 0.24 and 0.28 ng/μl for well-studied therapeutic antibodies like S309 and DXP-604, respectively. These values are 10~40-fold higher than those of the subset antibodies. Concerningly, some antibody drugs, such as REGN10933, REGN10987, LY-CoV555, LY-CoV016, AZD1061 and AZD8895, almost lost their neutralization activities against Omicron (Fig. 2b)^18^. Meanwhile, specific VOC-resistant antibodies with high neutralizing potency against WT and some other VOCs (IC_50_ <0.2 ng/μl) were identified and these comprise ~30% of the antibody repertoire, indicative of the evolution of a wide range of antibodies after 3-dose vaccination. Experiments repeated using authentic virus, including WT and all five VOCs, showed similar neutralization patterns by all these antibodies, further verifying the neutralizing potency and breadth for this subset of antibody repertoire elicited by 3-dose vaccination (Fig. 2e).

## Cryo-EM structures of the Omicron Spike in complex with 3 types of all five VOCs-reactive antibodies

Antibodies targeting the RBD can be categorized into six general classes (from I to VI) based on cluster analysis on epitope from 265 available RBD-NAb complex structures ^5^, that are related to the four groups on the basis of competition with the hACE2 for binding to S and recognition of the up or down state of the three RBDs in S ^19,20^. ELISA-based square competition matrix analysis with the aid of existing structural data revealed the presence of 3 major groups in this subset of antibody repertoire (Extended Data Fig. 2). To delineate the structural basis for antibody-mediated neutralization, we determined the cryo-EM structure of a prefusion stabilized Omicron S trimer in complex with representative Fab fragments. The two highly potent antibodies against Omicron (XGv347 and XGv289 with IC_50_ values of 0.006 and 0.016 ng/μl, respectively), one mAb (XGv282 with IC_50_ of 0.268 ng/μl) with median neutralizing activities against Omicron, but high neutralizing potency against other four VOCs, and one mAb (XGv265 with IC_50_ of 7.479 ng/μl) with >500-fold decreased neutralization against Omicron, but potent neutralization against other four VOCs were selected for structural investigations (Fig. 2b). We determined cryo-EM reconstructions of these complexes at 3.2 – 3.6 Å, and performed local refinement to further improve the densities around the binding interface between RBD and antibodies, enabling reliable analysis of the interaction details (Fig. 3, Extended Data Fig. 3, 4 and 5, Extended Data Table 1).

**Fig. 3.**
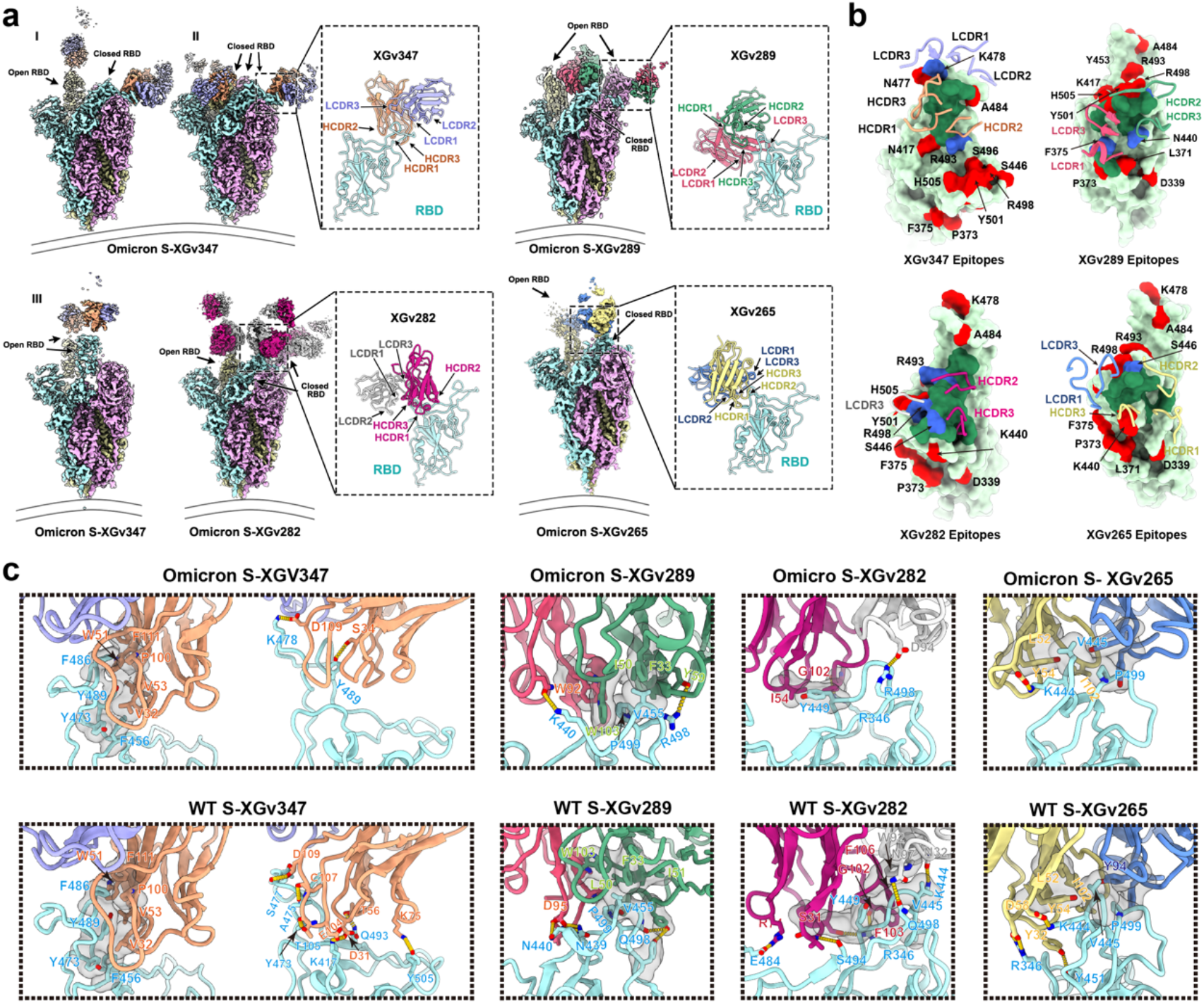
Structural basis of the broad and potent neutralization of representative antibodies. **a**. Cryo-EM maps and the binding modes of SARS-CoV-2 omicron S trimer in complex with XGv347 (top left), XGv289 (top right), XGv282 (bottom left) and XGv265 (bottom right). The three states of XGv347 binding to S-Omicron RBD are marked state I (one open and one closed RBD), state II (three closed RBDs) and state III (two open RBDs). Cartoon representations of the structure of SARS-CoV-2 Omicron-RBD in complex with the four antibodies are zoomed in. **b**. Interactions between the four antibodies and SARS-CoV-2 Omicron RBD. The CDRs of the four antibodies that interact with SARS-CoV-2 Omicron RBD are displayed as cartoon over the light green surface of the RBD. The mutation sites on RBD of Omicron are colored red, the epitopes of antibodies are colored deep green and the overlap of them are colored in blue. **c**. Interactions details between antibodies (XGv347, XGv289, XGv282 and XGv265) and SARS-CoV-2 Omicron (top) and WT RBD (bottom). All the WT structures are predicted with GROMACS. Hydrophobic patches and hydrogen bonds are highlighted with surface and dash lines. Color scheme is the same as in **a**.

The XGv347-Omicron S complex structures revealed three distinct conformational states: three XGv347 Fabs bound to a completely closed S with three down RBDs; two XGv347 Fabs bound to either two or one up and one down RBDs on S (Fig. 3a). By contrast, each of the complex structures for XGv289, XGv282 and XGv265 showed only one configuration where three XGv289 Fabs bound to two up and one down RBDs; three XGv282 Fabs bound to one up and two down RBDs; two XGv265 Fabs bound to S trimer with one down and one up RBD, although the XGv265-bound up RBD conformation was weakly resolved and therefore not modeled (Fig. 3a). Antibody XGv347 binds to an epitope at the tip of RBD, largely overlapping with the patch targeted by ACE2 (Fig. 2d, 3b, Extended Data Fig. 1). Structural comparisons revealed that XGv347 is very similar to A23-58.1, an ultrapotent and broadly reactive NAb effective against 23 SARS-CoV-2 variants ^21^. Furthermore, the residues of the epitope of XGv347 match with a major subset of those targeted by S2K146, another broadly cross-reactive sarbecovirus NAb ^22,23^, highlighting a plausible capability of these NAbs to cross-neutralize Omicron, SARS-CoV-2 variants and other sarbecoviruses through ACE2 molecular mimicry. Unexpectedly, the epitopes of XGv347, A23-58.1 as well as their sister NAbs would be normally inaccessible for the RBD-down conformation in the WT S, but become accessible for either up or down RBDs in the Omicron S due to a markedly outward expansion and a ~10° clockwise rotation of three RBDs, leading to an approximately 9 Å conformational movement for RBM (Fig. 3b and Extended Data Fig. 6). The XGv347 paratope constituted five complementarity determining regions (CDRs) with heavy chain and light chain contributing 70% and 30% of the binding surface area, respectively (Fig. 3b and Extended Data Table. 2). Overall XGv289, XGv282 and XGv265 bind patches surrounding the right shoulder of RBD with various orientations, but in a manner similar to those observed for DH1047, BD-812 and REGN10987; antibodies known to generally neutralize most VOCs with high potency ^24–26^, but showing declined, to varying degrees, binding and neutralizing activities against Omicron due to the presence of new N440K and G446S mutations (Fig. 2b, Extended Data Fig. 7 and Extended Data Table. 3). Notably, XGv265 and REGN10987 recognize almost same epitopes, both nearly losing their neutralizing activities against Omicron, despite retaining weak binding (Extended Data Fig. 7). Structural superimpositions reveal that XGv347 and either XGv289 or XGv265 can simultaneously bind to S, informing strategies to rationally design two-antibody cocktails (Extended Data Fig. 8).

## Structural basis for broad cross-neutralization activity of NAbs and immune escape

XGv347, XGv289, XGv282 and XGv265 bound Omicron with 5-40 folds lower affinity compared to their binding with WT, although the same binding modes for two orthologs were observed (Fig. 3a). For XGv347, tight binding to WT S is primarily due to extensive hydrophobic interactions contributed by F456, Y473, F486 and Y489 from WT RBD, V32, V53, W51, P100 and F111 from heavy chain, and Y33 from light chain, and 9 hydrogen bonds (Fig. 3c and Extended Data Table 3). Hydrophobic interactions between the Omicron RBD and XGv347 are largely maintained. However, substitutions of Y505H and K417N abolish three hydrogen bonds forged with K75, D31 and E104 from HCDRs, leading to conformational shifts in HCDR3 and the RBM tip (residues 470-490), which further perturb six hydrogen bonds built by Y473, A475, S477, T478, Q493 from WT RBD with T105, C107, A56, G55 and D109 from HCDRs, albeit with an extra hydrogen bond established by the mutation Q493R and G55 from HCDR2 in Omicron (Fig. 3c). Similarly, a large patch of hydrophobic interactions constructed by V445, G446, Y449, P499 from WT RBD and F33, L50, I51, Y59, W103 from HCDRs as well as extensive hydrophilic interactions facilitate tight binding between XGv289 and WT S (Fig. 3c). Substitution of G446S disrupts the hydrophobic microenvironment, substantially decreasing hydrophobic interactions between Omicron S and XGv289. Furthermore, mutations of N440K and Q498R, together with altered local conformation, also lessen hydrogen bonds formed by N439, K440, Y449, R498, T500, Q506 from Omicron RBD and D95, L98 from LCDRs as well as Y59, N62 from HCDRs that would exist in XGv289-WT S complex (Fig. 3c). Among these four representative antibodies, XGv282 showed minimal reduction in binding affinity (5-fold), but remarkable reduction in neutralization (~40-fold), versus the characterization of XGv347 with 40-fold decrease in binding, but unchanged neutralization against Omicron when compared to WT, suggesting that epitope, rather than binding affinity, might play more crucial roles in the neutralizing potency and breadth of an antibody. Consistent with XGv289, the substitution of G446S alters the hydrophobic microenvironment generally established by RBD and a group of antibodies bound at the right shoulder, including XGv289 and XGv282, triggering a conformational shift on CDRs and disrupting antibody recognition (Fig. 3c). In addition, the mutation E484A breaks hydrogen bond-connection with R74 from XGv282 HCDR2 and losses of charge interactions between R346, K444 from WT RBD and D56, D58 of XGv265 LCDR2 due to conformational alterations, further decreasing the binding of XGv282 and XGv265 to the Omicron variant, respectively (Fig. 3c). Taken together, G446S, acting as a critical mutation site, can alter the local conformation at the binding interface, conferring greater resistance to one class of antibodies bound at the right shoulder of RBD.

## The therapeutic activities of five representative antibodies against the Beta SARS-CoV-2 variant in mice

Given the excellent neutralizing breadth and potency at cell-based levels for above antibodies, we next sought to assess the correlation between *in vitro* neutralization and *in vivo* protection. A number of representative mAbs with high neutralizing potency and breadth, belonging to different classes, such as XGv347, XGv289, XGv282, XGv265 and XGv052, produced in the HEK293F cell line were selected for therapeutic evaluation in a well-established mouse model challenged with the Beta variant ^27^. Upon Beta intranasal challenge, adult BALB/c showed robust viral replication in the lungs at 3-5 days post inoculation. To evaluate the protection efficacy of these mAb, BALB/c mice challenged with the Beta variant were administered a single dose of as low as 5 mg/kg of XGv347, XGv289, XGv282, XGv265 and XGv052 individually or combinations of XGv282 and XGv347 (2.5 mg/kg for each), and XGv052 and XGv289 (2.5 mg/kg for each) in therapeutic settings (Fig. 4a). Heavy viral loads with high levels of viral RNAs (> 10^9^ copies/g) were detected in the lungs at 5 day post infection in the control group of mice treated with PBS. However, a single dose of XGv282 reduced the viral RNA loads by ~10,000-fold in the lungs compared to the control group (Fig. 4b). Remarkably, a single dose of XGv289, XGv265, XGv347, XGv052 or antibody cocktails of XGv282 + XGv347, XGv052 + XGv289 resulted in a complete clearance of viral particles in the lungs (Fig. 4b and 4c). A potential synergistic effect was observed for combined therapies of XGv282 + XGv347 at 2.5 mg/kg for each (Fig. 4b and 4c). In addition, histopathological examination revealed severe interstitial pneumonia, characterized by alveolar septal thickening, inflammatory cell infiltration and distinctive vascular system injury developed in mice belonging to the control group at day 5 (Fig. 4d). In contrast, no obvious lesions of alveolar epithelial cells or focal hemorrhage were observed in the lung sections from mice that received indicated antibody treatments (Fig. 4d). Collectively, these results suggest that some of the antibodies from the repertoire elicited by a 3-dose vaccination regimen retain therapeutic potential against all circulating VOCs; albeit the protective efficacy against Omicron needs to be investigated more thoroughly.

**Fig. 4.**
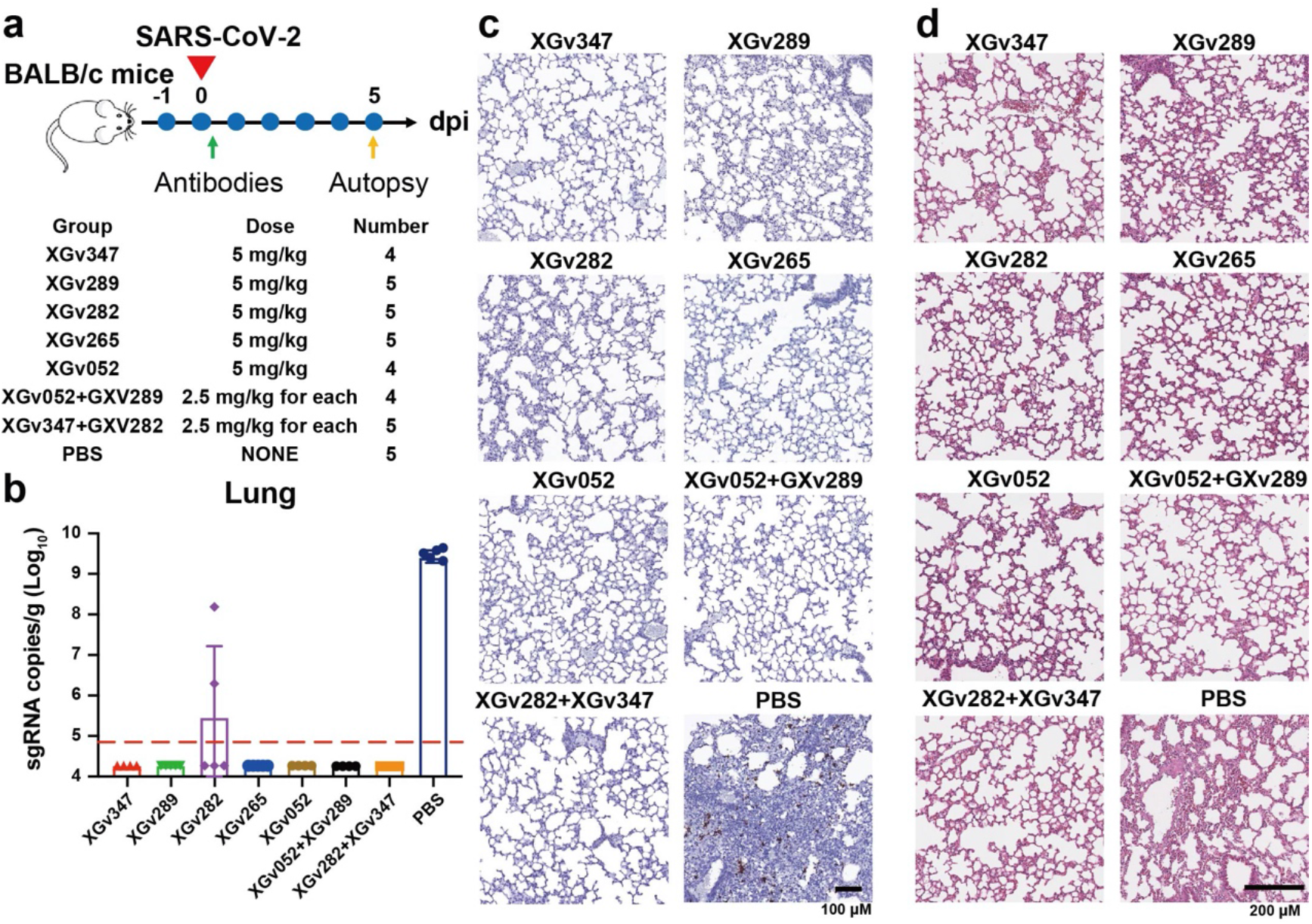
Protection against SARS-CoV-2 Beta variant strain challenge in mice. **a,** Experimental design. Groups of BALB/c mice were infected intranasally with SARS-CoV-2 Beta variant strain, followed by a single dose of an antibody, or an antibody cocktail, or PBS as control one hour after infection as indicated in **a**. **b to d**, Examination of lung tissues of mice collected at 5 dpi for **b**, virus titer, **c**, Immunostaining and **d**, H&E. **b**, Virus RNA loads in the lungs at 5 dpi were measured by RT-qPCR and are expressed as RNA copies per gram. Data are represented as mean ± SD. Dashed line represents limit of detection. **c**, SARS-CoV-2 genome RNA ISH was performed with a SARS-CoV-2 specific probe. Brown-colored staining indicates positive results. Scale bar, 200 μm. **d**, Histopathological analysis of lung samples at 5 dpi. Scale bar: 100 μm.

## Discussion

The ongoing pandemic has witnessed frequent occurrences of SARS-CoV-2 variants that increase transmissibility and reduce potency of vaccine-induced and therapeutic antibodies ^2,11^. More recently, there has been unprecedented concern that the Omicron variant has significantly increased antibody escape breadth due to newly occurred and accumulated mutations in the key epitopes of most neutralizing antibodies. Alarmingly, Omicron nearly ablates the neutralization activity of most FDA approved antibody drugs, including LY-CoV555, LY-CoV016, REGN10933, REGN10987, AZD8895 and AZD1061 ^18^. These issues raise an urgent need to develop next-generation antibody-based therapeutics that can broadly neutralize these variants, as well as future variants of concern. Our previous study revealed that the regimen of 3-dose vaccination (0, 1, 7 months) of inactivated vaccine leads to an improved immunity response with significantly enhanced neutralizing breadth via ongoing antibody somatic mutation and memory B cell clonal turnover ^5,28^. Correlated with this, one subset of highly potent neutralizing antibodies with broad activities (IC_50_ < 0.2 ng/μl) against all circulating VOCs, including Omicron, were present in at least four individuals who had received three doses of inactivated ancestral SARS-CoV-2 vaccine. Some, but not limited to these of this subset antibodies fully protected against the Beta variant infection in mice, although their *in vivo* breadth and protective efficacy against Omicron remains unconfirmed. Furthermore, our structural and functional analyses revealed that a newly occurred mutation, G446S, might act as a critical antibody escape site, conferring greater resistance to one major class of antibodies bound at the right shoulder of RBD via altering microenverionments at the S-NAb binding interface.

In addition to evading currently available antibody therapeutics, the Omicron variant can diminish the efficacy of all clinically approved vaccines, including the mRNA vaccines and inactivated vaccines ^11,12^. There is an ongoing debate about whether the immune responses can be fine-turned to the Omicron variant by boosting with a tweaked (Omicron-based) vaccine. A major hurdle for this approach is the “original antigenic sin”, a phenomenon documented in some other infectious diseases, including flu ^29^. The presence of a subset of antibodies with broad neutralizing activities against all circulating VOCs in memory B-derived antibody repertoire from the 3-dose vaccinees suggests a possibility that selective and expeditious recall of humoral responses might be elicited via the Omicron/future variants infection, conferring to a secondary protection directed by memory etched in the immune system. Further studies are warranted to examine the advantages and disadvantages of booster shots of an Omicron-specific vaccine or simply administration of a booster with the original vaccines. Lastly the identification and characterization of broadly protective antibodies against all circulating VOCs will aid in the development of universal vaccination strategies against sarbecoviruses.

## Materials and Methods

### Facility and ethics statements

All procedures associated with SARS-CoV-2 live virus were approved by the Animal experiment Committee Laboratory Animal Center, Beijing Institute of Microbiology and Epidemiology with an approval number of IACUC-IME-2021-022 and performed in Biosafety Level 3 (BSL-3) laboratories in strict accordance with the recommendations in the Guide for Care and Use of Laboratory Animals.

### Viral stock and cell lines

SARS-CoV-2 wild-type strain CN01 was originally isolated from a patient during the early phase of COVID-19 endemic in China. SARS-CoV-2 variant of concern (VOC) Beta (B.1.351 lineage) strain GDPCC was isolated in a patient from South Africa and an Omicron (B.1.1.529 lineage) strain was isolated from a patient in Hong Kong and now preserved in SinoVac Biotech Ltd. All virus strains were first purified by standard plaque assay as previously described ^13^ and then inoculated into Vero cells (CCL-81) grown to 95% in 10% fetal bovine serum (FBS) supplemented Dulbecco’s minimal essential medium (DMEM) for amplification.

### Human sera samples

The serum samples were obtained from healthy volunteers who had no history of COVID-19 and were verified by PCR and serological assay and received two doses or three doses of CoronaVac (Sinovac) inactivated vaccine specific against SARS-COV-2. All volunteers were provided informed written consent form and the whole study was conducted in accordance with the requirements of Good Clinical Practice of China.

### Authentic virus neutralization assay

The serum samples were first incubated at 56 °C for 30 min for inactivation. The heat-treated samples or monoclonal antibodies (mAbs) were subject to seral dilution from 1: 4 or 50 ng/μL with DMEM in two-fold steps and mixed with a virus suspension containing 100 TCID_50_ at 36.5°C for 2h, after which, the mixtures were added to wells seeded with confluence Vero cells and incubated at 36.5°C for another 5 days in a humidified 5% CO_2_ cell incubator. After that, the cytopathic effect (CPE) of each well was observed under microscopes by three different individuals and the related dilutions and concentrations were recorded and used for the titration of samples tested by the method of Reed-Muench ^13^.

### Pseudovirus neutralization assay

The pseudotyped viruses bearing the Spike (S) protein were generated, aliquoted and restored as previously described ^17^. Briefly, 293T cells were first transfected with the plasmid embedded with the S gene of wild-type or VOC/VOI (Alpha, Beta, Gamma, Delta, Lamda and Omicron) SARS-CoV-2. The transfected 293T cells were infected with VSV G pseudotyped virus (G*ΔG-VSV) at a multiplicity of infection (MOI) of 4. After incubation for five hours, cells were washed with PBS, and then complete culture medium was added. After another 24 hours, the SARS-CoV-2 pseudoviruses were produced and harvested. For the *In vitro* pseudotyped virus neutralization assay, the plasma samples or antibodies were diluted in DMEM starting from 1:10 or 10 ng/μL with 6 additional threefold serial dilutions, each of which were mixed with the harvested pseudovirus and incubated at 37 °C for 1h. After that, the mixtures were added to Huh-7 cells and placed back for incubation for another 24 hours. Then, the luciferase luminescence (RLU) of each well was measured with a luminescence microplate reader. The neutralization percentage was calculated as following: Inhibition (%) = [1- (sample RLU-Blank RLU) / (Positive Control RLU-Blank RLU)] (%). Antibody neutralization titers were presented as 50% maximal inhibitory concentration (IC_50_).

### Protein expression and purification

The sequences of VOC Omicron full-length spike (S) protein (residues 1-1208), receptor-binding domain (RBD) (residues 319-541) and N-terminal domain (NTD) (residues 1-304) were modified from the plasmids encoding the S, RBD and NTD of wild-type SARS-COV-2 (GenBank: MN908947) in our lab by overlapping PCR. In additional to the reported mutations (A67V, Δ69-70, T95I, G142D, Δ143-145, Δ211, L212I, ins214EPE, G339D, S371L, S373P, S375F, K417N, N440K, G446S, S477N, T478K, E484A, Q493K, G496S, Q498R, N501Y, Y505H, T547K, D614G, H655Y, N679K, P681H, N764K, D796Y, N856K, Q954H, N969K, L981F) on Omicron, the proline substitutions at 986 and 987, ‘GSAS’ substitutions at the S1/S2 furin cleavage site (residues 682-685) and a C-terminal T4 foldon trimerization domain were also remained in the Omicron S construct to stabilize the trimeric conformation of S protein. For protein expression, the plasmids of these proteins were transiently transfected into HEK293 F cells grown in suspension at 37 °C in an incubator supplied with 8% CO_2_, rotating at 130 rpm. The cell supernatants were harvested and concentrated three days post-transfection, and further purified by affinity chromatography using resin attached with streptavidin or Ni-NTA and size-exclusion chromatography (SEC) using a Superdex 200 column (GE Healthcare Life Sciences) with the buffer containing 20 mM Tris pH 8.0 and 200 mM NaCl.

### Antibody expression and Fab generation

The selected 323 antibodies were subjected to gene codon optimization, construction and expression as described previously ^5^. Then the clones were transiently transfected into mammalian HEK293F cells and incubated for 5 days in a 5% CO_2_ rotating incubator at 37°C for antibody expression, which were further purified using protein A. The purified mAbs XGv265, XGv282, XGv289 and XGv347 were then processed to obtain their Fab fragments using the Pierce FAB preparation kit (Thermo Scientific) as described previously ^30^. Briefly, the samples were first applied to desalination columns to remove the salt and the flow-throughs were collected and incubated with papain that was attached with beads to cleave Fab fragments from the whole antibodies for 5 hours at 37°C. After that, the mixtures were transferred into Protein A columns and the flow-throughs, i.e., the Fab fragments were collected and dialyzed into Phosphate Buffered Saline (PBS) (ThermoFisher, catalog #10010023).

### Bio-layer interferometry

Bio-layer interferometry (BLI) experiments were run on an Octet Red 384 machine (Fortebio). To measure the binding affinities of mAbs, monoclonal antibodies were immobilized onto Protein A biosensors (Fortebio) and the threefold serial dilutions of wild-type RBD (ACROBiosystems, Cat No. SPD-C52H3), Alpha RBD (ACROBiosystems, Cat No. SPD-C52Hn), Beta RBD (ACROBiosystems, Cat No. SPD-C52Hp), Gamma RBD (ACROBiosystems, Cat No. SPD-C52Hr), Delta RBD (ACROBiosystems, Cat No. SPD-C52Hh) and Omicron RBD (ACROBiosystems, Cat No. SPD-C522e) were used as analytes. Data were then analyzed using software Octet BLI Analysis 12.2 (Fortebio) with a 1:1 fitting model. For the epitope binning by BLI, SARS-CoV-2 VOC Omicron RBD tagged with his (ACROBiosystems, Cat No. SPD-C522e) was loaded on HIS1K biosensors, which were pre-equilibrated in the buffer for at least 1 min. The loaded biosensors were immersed with the first mAb for 300 s, followed by addition of the second mAb for another 90 s. Date obtained were also analyzed by Octet BLI Analsis 12.2.

### ELISA assays

To evaluate whether the given mAbs could block the interaction between human ACE2 (hACE2) and RBD, ACE2 competition ELISA was performed by using the SARS-CoV-2 (B.1.1.529) Inhibitor Screening Kit (ACROBiosystems, Cat No. EP-115) according to the recommended protocol. Briefly, each of the 10 two-fold dilution series of mAbs (starting dilution of 25 ng/μL) and 0.8 ng/μL of HRP-conjugated SARS-CoV-2 Omicron RBD were added into the ELISA plate wells which are pre-coated with hACE2 protein. After incubation at 37 °C for 1 hour, the plates were washed three times with PBST (0.1% Tween) and the colorimetric signals were developed by addition of 3, 3’, 5, 5’-tetramethylbenzidine TMB (Thermo Fisher) for 10 min. The reaction was stopped by addition of 50 μL of 1M H2SO4. The absorbance was measured at 450 nm with an ELISA microplate reader. For each mAb, a blank control with no mAb was added for inhibition calculation. The area under the curve (AUC) of each mAb were determined using Prism V8.0 (GraphPad). For competitive ELISAs to identify the domain of a given mAb, 96-well plates were first coated with RBD (2 μg/ml) and then blocked with 2% BSA in PBS. After incubation with the reference mAbs, the blocking antibody (15 μg/ml), the wells were followed by directly adding the second biotinylated antibodies (0.25 μg/ml). Streptavidin-HRP (BD Biosciences) was then added for detection. Samples with no first antibody were used as a negative control for normalization.

### Cryo-EM sample preparation, data collection

The purified S protein was mixed with each of the Fab fragments of XGv265, XGv282, XGv289 or XGv347 with a molar ratio of 1: 1.2 for 10 s ice incubation, and then dropped onto the pre-glow-discharged holey carbon-coated gold grid (C-flat, 300-mesh, 1.2/1.3, Protochips In.), blotted for 7 seconds with no force in 100% relative humidity and immediately plunged into the liquid ethane using Vitrobot (FEI). Cryo-EM data sets of these complexes were collected at 300 kV with an FEI Titan Krios microscope (FEI). Movies (32 frames, each 0.2 s, total dose of 60 e^-^ Å^−2^) were recorded using a K3 Summit direct detector with a defocus range between 1.5-2.7 μm. Automated single particle data acquisition was carried out by SerialEM, with a calibrated magnification of 22,500 yielding a final pixel size of 1.07 Å.

### Cryo-EM data processing

A total of 3,752, 2,631, 3,955 and 5,014 micrographs of S-XGv265-complex, S-XGv282-complex, S-XGv289-complex and S-XGv347-complex, respectively were recorded and subjected to beam-induced motion correction using motionCorr in Relion3.0 package ^31^. The defocus value of each image was calculated by Gctf. Then, 1,302,103, 756,508, 2,332,045 and 2,320,416 particles of the S-XGv265-complex, S-XGv282-complex, S-XGv289-complex and S-XGv347-complex, respectively were picked and extracted for reference-free 2D alignment by cryoSPARC ^32^, based of which, 422,083, 190,154, 837,832 and 614,852 particles were selected and applied for 3D classification by Relion3.0 for S-XGv265-complex, S-XGv282-complex, S-XGv289-complex and S-XGv347-complex, respectively with no symmetry imposed to produce the potential conformations for the complexes. Afterwards, the candidate model for each complex was selected and processed by non-uniform auto-refinement and postprocessing in cryoSPARC to generate the final cryo-EM density for S-XGv265-complex, S-XGv282-complex, S-XGv289-complex and S-XGv347-complex. To improve the resolution of the interface between RBD and mAbs, the block-based reconstruction was performed to obtain the final resolution of the focused interfaces which contained the interfaces of RBD and mAbs investigated here as described previously ^33^. The resolution of each structure was determined on the basis of the gold-standard Fourier shell correlation (threshold = 0.143) and evaluated by ResMap. All dataset processing is shown in Extended Data Fig. 3 and also summarized in Extended Data Table 1.

### Model fitting and refinement

The atomic models of the complexes were generated by first fitting the chains of the native apo SARS-CoV-2 S trimer (PDB number of 6VYB) and Fabs (PDB number of 7LSS and 7CZW for XGv265, 5MES and 5VAG for XGv282, 6UDA and 7MEG for XGv289 as well as 7E3K for XGv347) into the cyo-EM densities of the final S-Fab-complexes described above by Chimera, followed by manually adjustment and correction according to the protein sequences and densities in Coot, as well as real space refinement using Phenix. Details of the refinement statistics of the complexes are summarized in Extended Data Table 1.

### MD simulation and ΔG estimation

Model of SARS-CoV-2 wild-type RBD in complex with XGv265, XGv282, XGv289 and XGv347 were generated in Chimera by superimposition of WT RBD and cryoEM structure of Omicron RBD in complex with the four antibodies. Before molecular dynamics, all models were checked by WHAT IF Web Interface (https://swift.cmbi.umcn.nl/servers/html/index.html) to model missing sidechains and remove atomic clashes. After that, the structure was simulated by GROMACS-2021. Briefly, we used OPLS force field with TIP3P water model to prepare the dynamic system and add Na+ and Cl^-^ ions to make the system electrically neutralized. Then, the system was subjected to energy minimization using the steepest descent algorithm until the maximum force of 1,000 kJ mol-1 has been achieved. NVT ensemb1e via the Nose-Hoover method at 300 K and NPT ensemble at 1 bar with the Parinello-Rahman algorithm were employed successively to make the temperature and the pressure equilibrated, respectively. Finally, a MD production runs of 100 ns were performed starting from random initial velocities and applying periodic boundary conditions. The non-bonded interactions were treated using Verlet cut-off scheme, while the long-range electrostatic interactions were treated using particle mesh Ewald (PME) method. The short-range electrostatic and van der Waals interactions were calculated with a cut-off of 12 Å. Average structure of the four complexes were generated using the last 10 ns frames and ΔG between the antibodies and RBD was estimated in ROSETTA by InterfaceAnalyzer. Atomic_burial_cutoff, sasa_calculator_probe_radius and interfaces_cutoff values were set to 0.01, 1.4 and 8.0 respectively.

### Protection against SARS-CoV-2 Beta variant strain challenge in mice

The in vivo protection efficacies of single antibody or antibody cocktail were assessed by using a newly established mouse model based on a SARS-CoV-2 Beta variant strain ^27^. Briefly, group of 8-month-old female BALB/c mice were infected with 1×10^4^ PFU of SARS-CoV-2 Beta variant strain, then infected mice were treated intraperitoneally with a single dose of different antibodies or antibody cocktails at 1 hour after infection. The lung tissues of mice were collected at 5 dpi for viral RNA loads assay and pathological examination.

### Viral burden determination

Viral burden in lung from mice were measured as described previously ^16^. Briefly, lung tissue homogenates were clarified by centrifugation and viral RNA was extracted using the QIAamp Viral RNA Mini Kit (Qiagen). Viral sgRNA quantification in each tissue sample was performed by quantitative reverse transcription PCR (RT-qPCR) targeting the S gene of SARS-CoV-2. RT-qPCR was performed using One-Step PrimeScript RT-PCR Kit (Takara).

### Histology, and RNA in situ hybridization (RNA ISH)

Lung tissues from mice were fixed with perfusion fixative (formaldehyde) for 48 h, and embedded in paraffin according to standard histological assays. For histopathology, lung tissues were stained with hematoxylin and eosin (H&E). Images were captured using Olympus BX51 microscope equipped with a DP72 camera. For RNA ISH assays were performed with an RNAscope 2.5 (Advanced Cell Diagnostics) according to the manufacturer’s instruction. Briefly, formalin-fixed paraffin-embedded tissue sections of 5 μm were deparaffinized by incubation for 60 min at 60 °C. Endogenous peroxidases were quenched with hydrogen peroxide for 10 min at room temperature. Slides were then boiled for 15 min in RNAscope Target Retrieval Reagents and incubated for 30 min in RNAscope Protease Plus before probe hybridization. The probe targeting 2019-nCoV RNA was designed and synthesized by Advanced Cell Diagnostics (catalog no. 848561). Tissues were counterstained with Gill’s hematoxylin and visualized with standard bright-field microscopy. Original magnification was 10×.

## Reporting summary

Further information on research design is available in the Nature Researh Reporting Summary linked to this paper.

## Data availability

The atomic coordinates of XGv347 in complex with S trimer (state I), XGv347 in complex with S trimer (state II), XGv347 in complex with S trimer (state III), XGv347-S have been submitted to the Protein Data Bank with accession numbers: 7WEA, 7WEC and 7WEB, respectively. Futhermore, the atomic coordinates of XGv265, XGv282 and XGv289 have been deposited in the protein data bank under accession code 7WE8, 7WE7 and 7WE9, respectively. Cryo-EM density maps in this study have been deposited at the Electron Microscopy Data Bank with accession codes EMD-32444 (state1), EMD-32446 (state2) and EMD-32445 (state3), EMD-32441 (XGv282), EMD-32442 (XGv265), and EMD-32443 (XGv289). To reveal structural details of Fab binding mechanism, the local optimized method are used to optimized data progress and the related atomic models and EM density maps of optimized reconstructions of Fab interaction interface has been deposited under accession code 7WEE (XGv265), 7WED (XGv347), 7WEF (XGv289), EMD-32447 (XGv347), EMD-32448 (XGv265), EMD-32449 (XGv289), respectively.

## Acknowledgments

We thank Dr. Xujing Li, Dr. Xiaojun Huang and Lihong Chen for cryo-EM data collection at the Center for Biological imaging (CBI) in Institute of Biophysics for EM work. We also thank Dr. Yuanyuan Chen, Zhenwei Yang, Bingxue Zhou for technical support on SPR. Work was supported by the Strategic Priority Research Program (XDB29010000, XDB37030000), CAS (YSBR-010), National Key Research and Development Program (2020YFA0707500, 2018YFA0900801), Beijing Municipal Science and Technology Project (Z201100005420017) and Ministry of Science and Technology of China (EKPG21-09 and CPL-1233). Xiangxi Wang was supported by Ten Thousand Talent Program and the NSFS Innovative Research Group (No. 81921005). Kang Wang was supported by the Special Research Assistant Project of the Chinese Academy of Sciences.

## Author contributions

X.W., K.W., C.F.Q., C.Q., and Y.W designed the whole study; K.W., Z.J., L.B., L.C., H.C., Y.H., Q.L., Y.J., Q.Z., Y.D., L.W., M.L., Y.L., K.F., P.Y., X.P., Z.C., L.Q., P.G., J.W., S.L., Y.C., W.H. performed experiments; K.W., L.W., P.L., N.W., W.F. H.C. prepared the Cryo-EM samples and solved the structures; all authors analyzed data; X.W., K.W., C-F.Q, C.Q., and Y.W. wrote the manuscript with input from all co-authors.

## Competing interests

All authors have no competing interests.

## Extended Data

**Extended Data Fig. 1.**
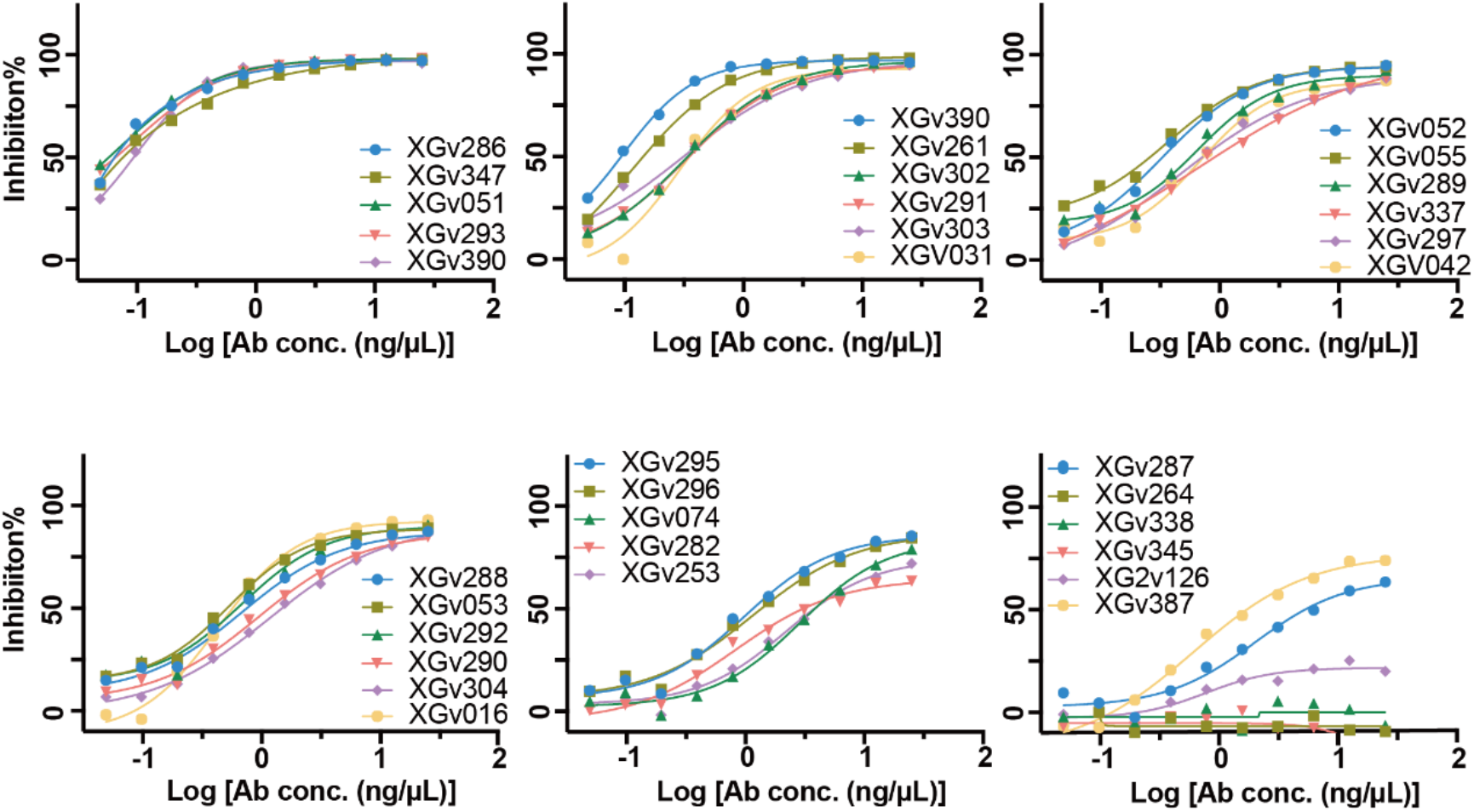
Fab-ACE2 Competition ELISA assay. Data shown are the curves of 31 antibodies used to compete with ACE2

**Extended Data Fig. 2.**
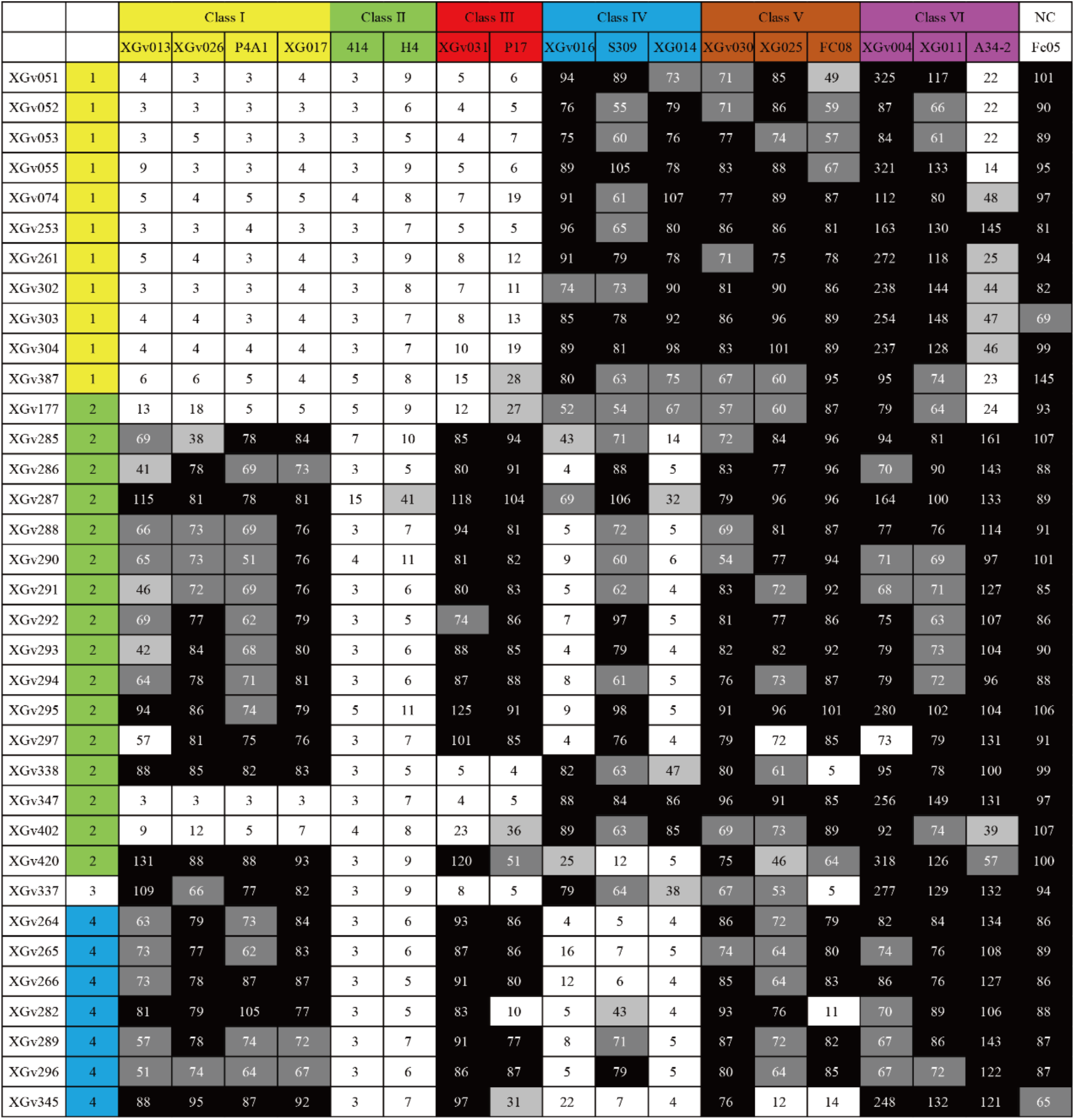
Data Sheets of ELISA assay of representative Mabs against Omicron RBD. Different Classes of Nabs (Class I-VI) are colored by yellow, green, red, blue, brown and magenta, respectively. Values are filled with black (>75), grey (50-75), silver (25-50) and white (<25).

**Extended Data Fig. 3.**
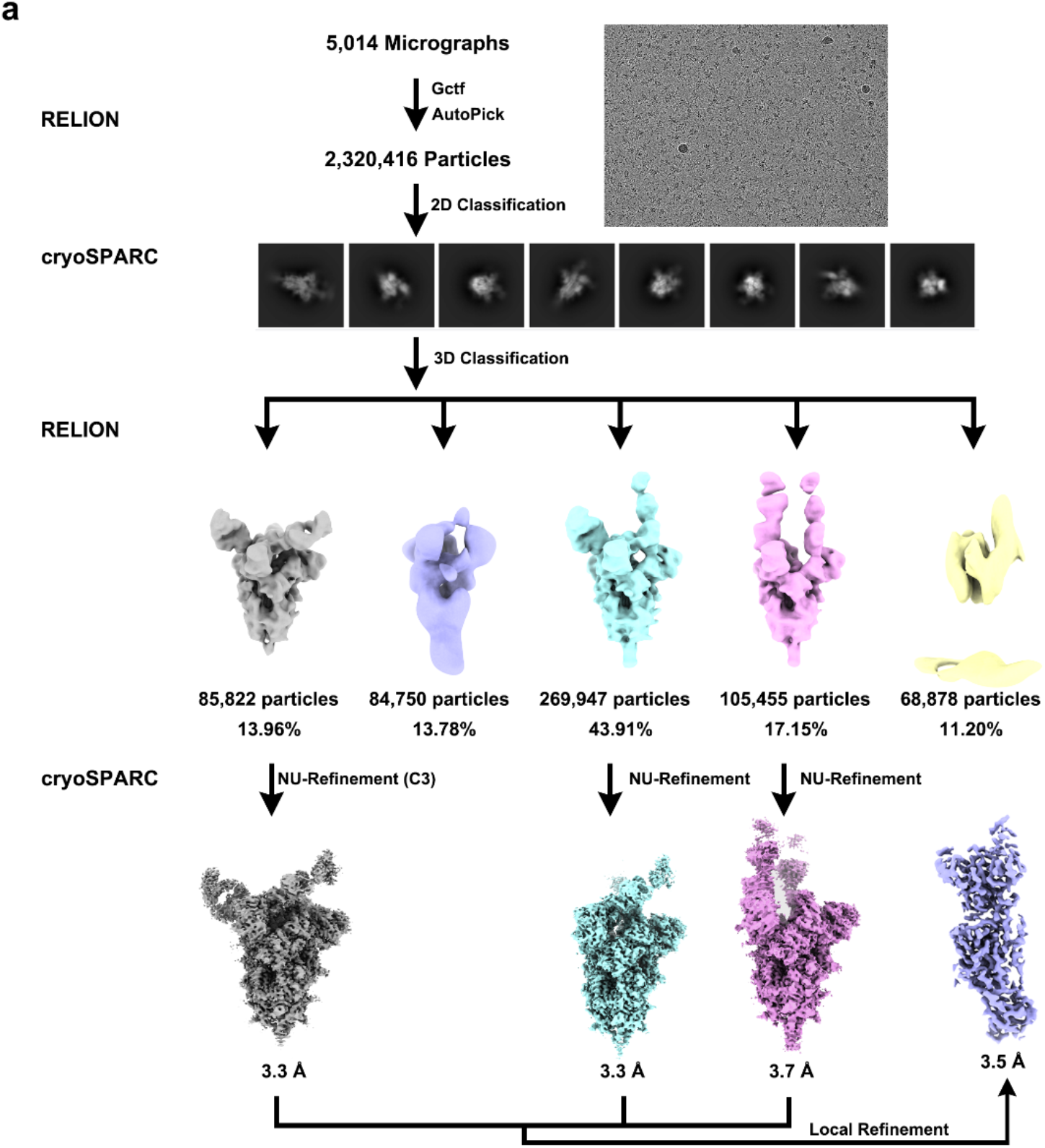
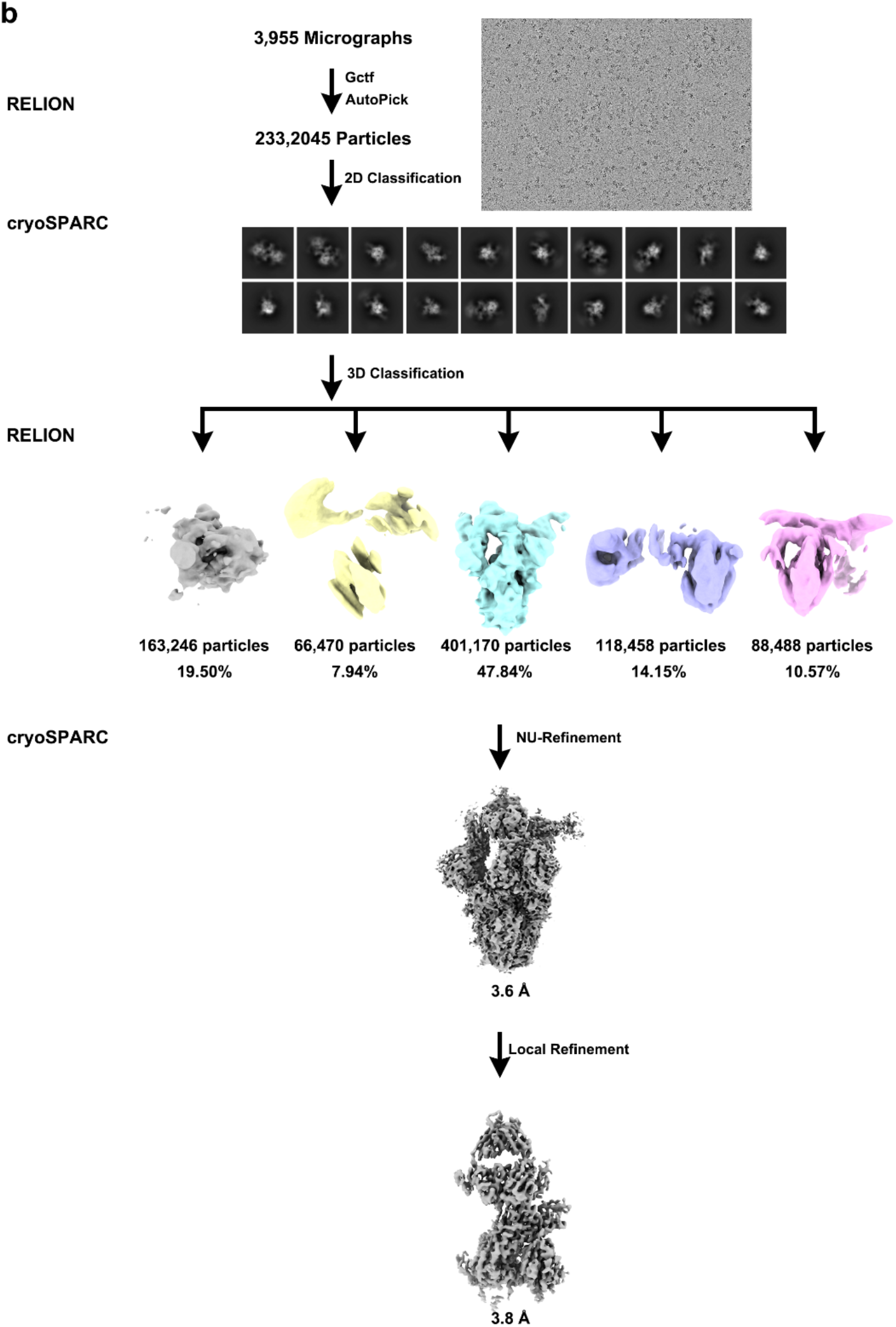
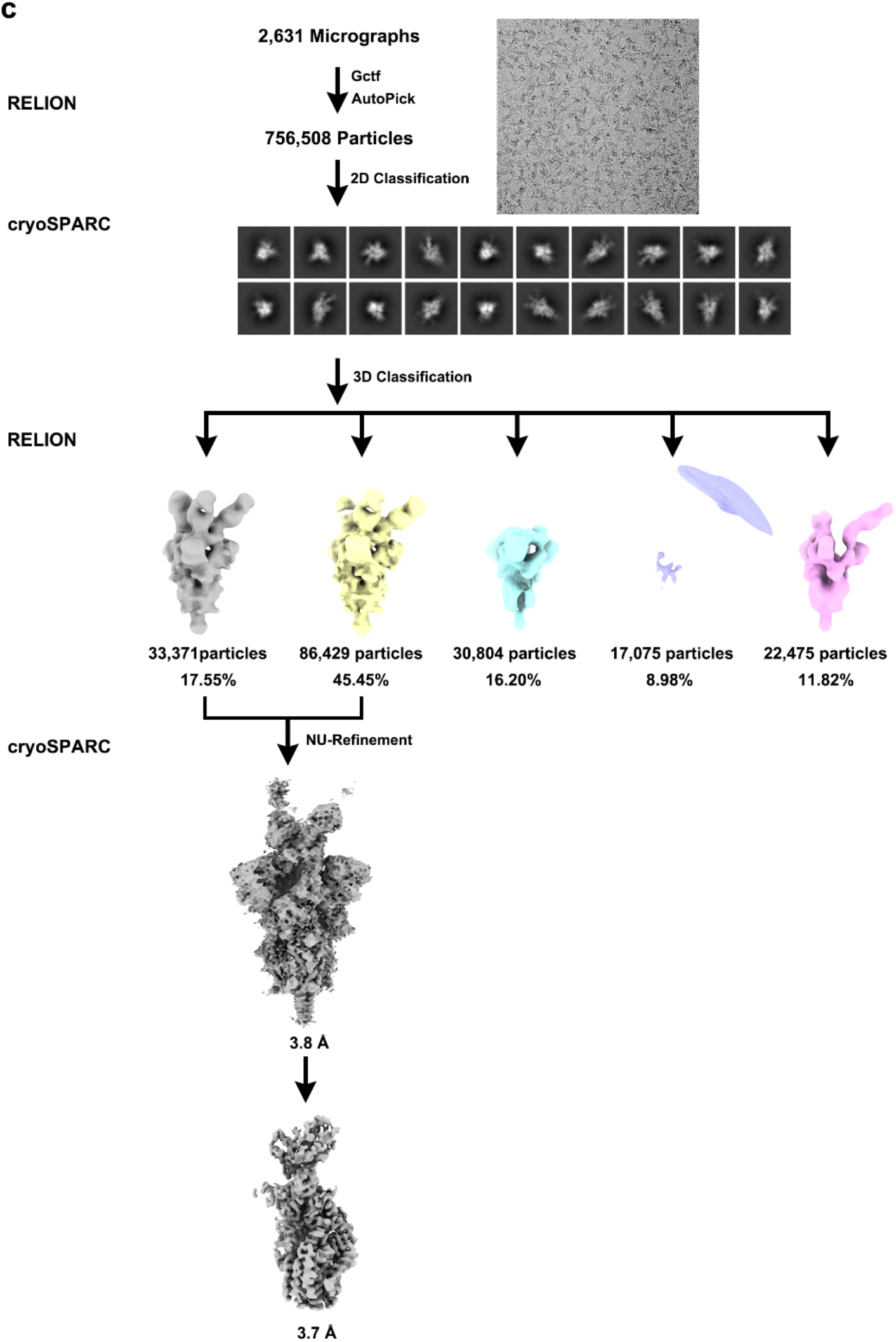
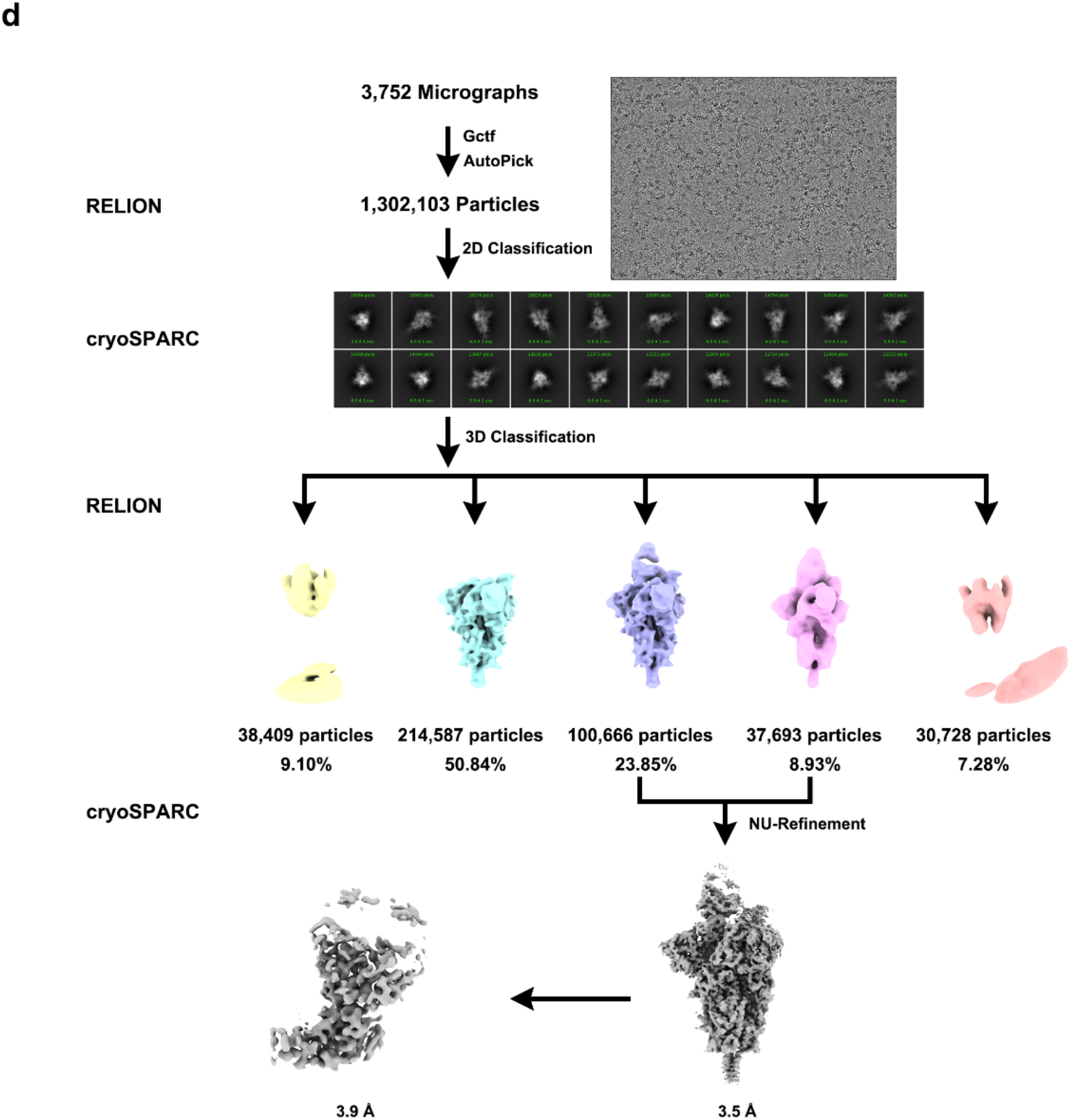
Flowcharts for cryo-EM data processing. Flowcharts for Omicron Spike protein in complex with **a**, Fab XGv347, **b**, XGv289, **c**, XGv282 and **d**, XGv265 are shown.

**Extended Data Fig. 4.**
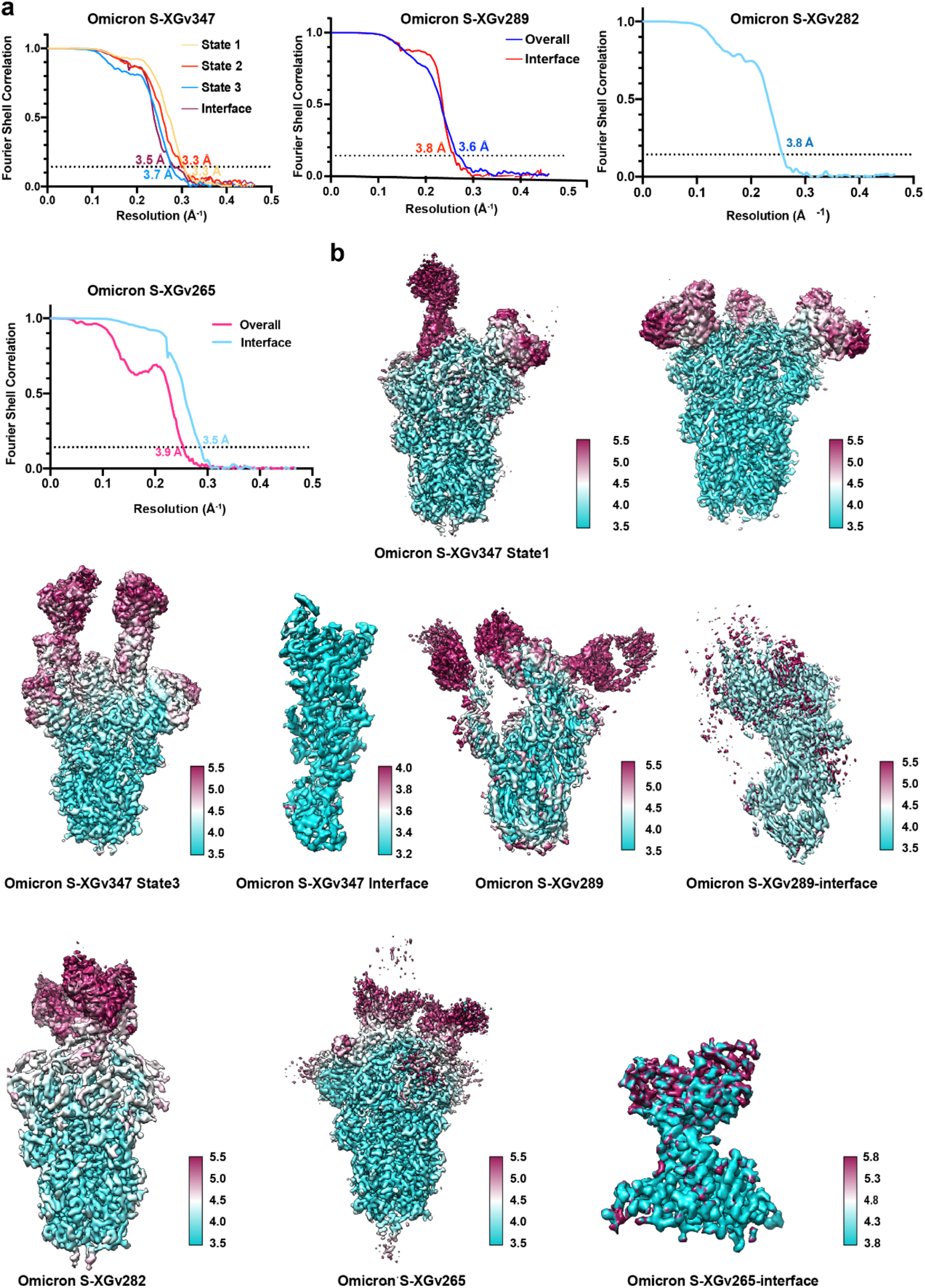
Resolution Estimation of the EM maps. **a,** The gold-standard FSC curves of overall maps of Omicron S trimer in complex with Fab XGv347, XGv289, XGv282 and XGv265 and local maps of interfaces. **b,** Local resolution assessments of cryo-EM maps using ResMap are shown.

**Extended Data Fig. 5.**
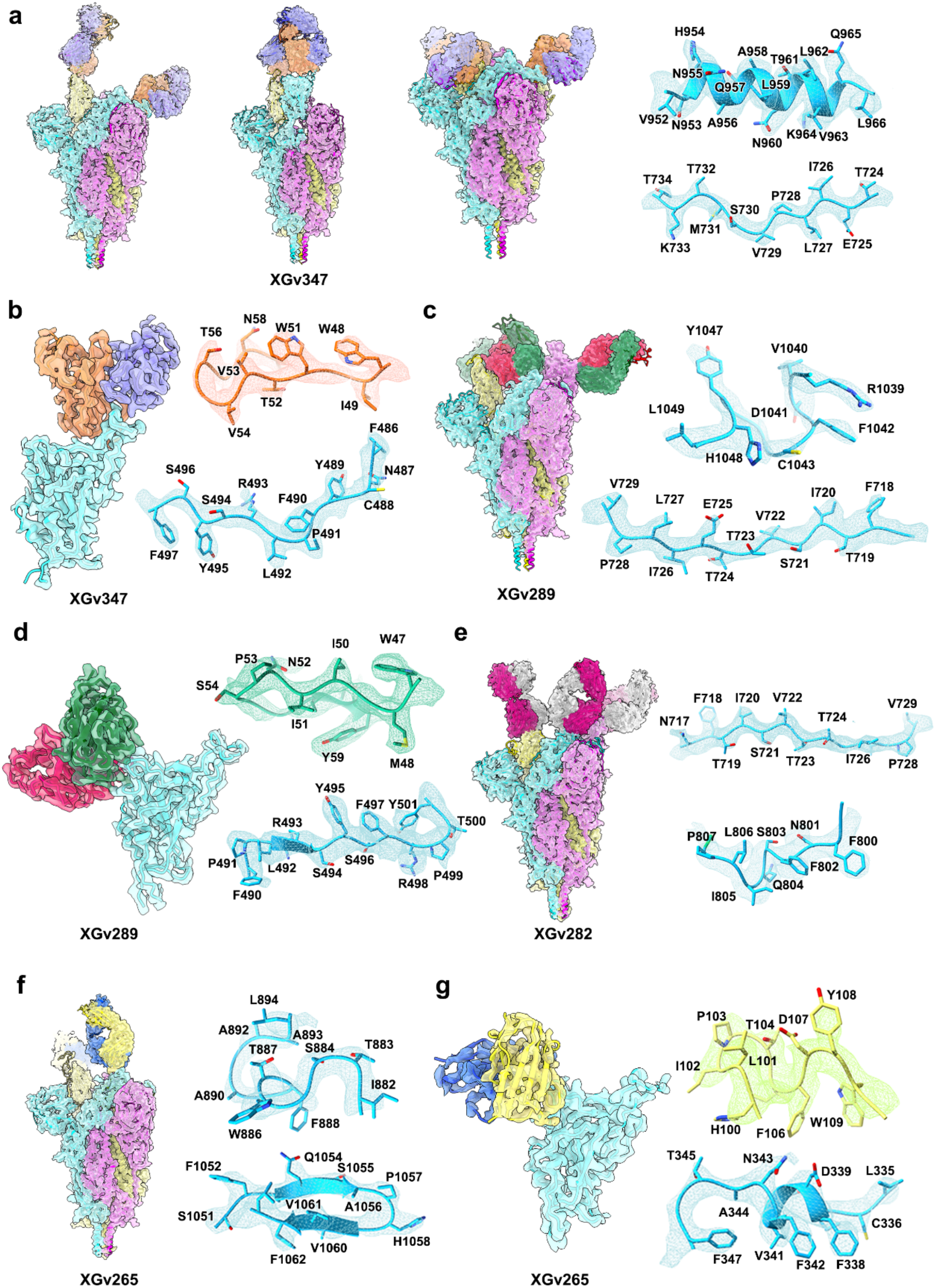
Density maps and atomics models. Cryo-EM maps of Omicron S trimer in complex with Fab XGv347, XGv289, XGv282 and XGv265 and their interfaces are shown. Color scheme is the same as in Fig. 3a. Residues are shown as sticks with oxygen colored in red, nitrogen colored in blue and sulfurs colored in yellow.

**Extended Data Fig. 6.**
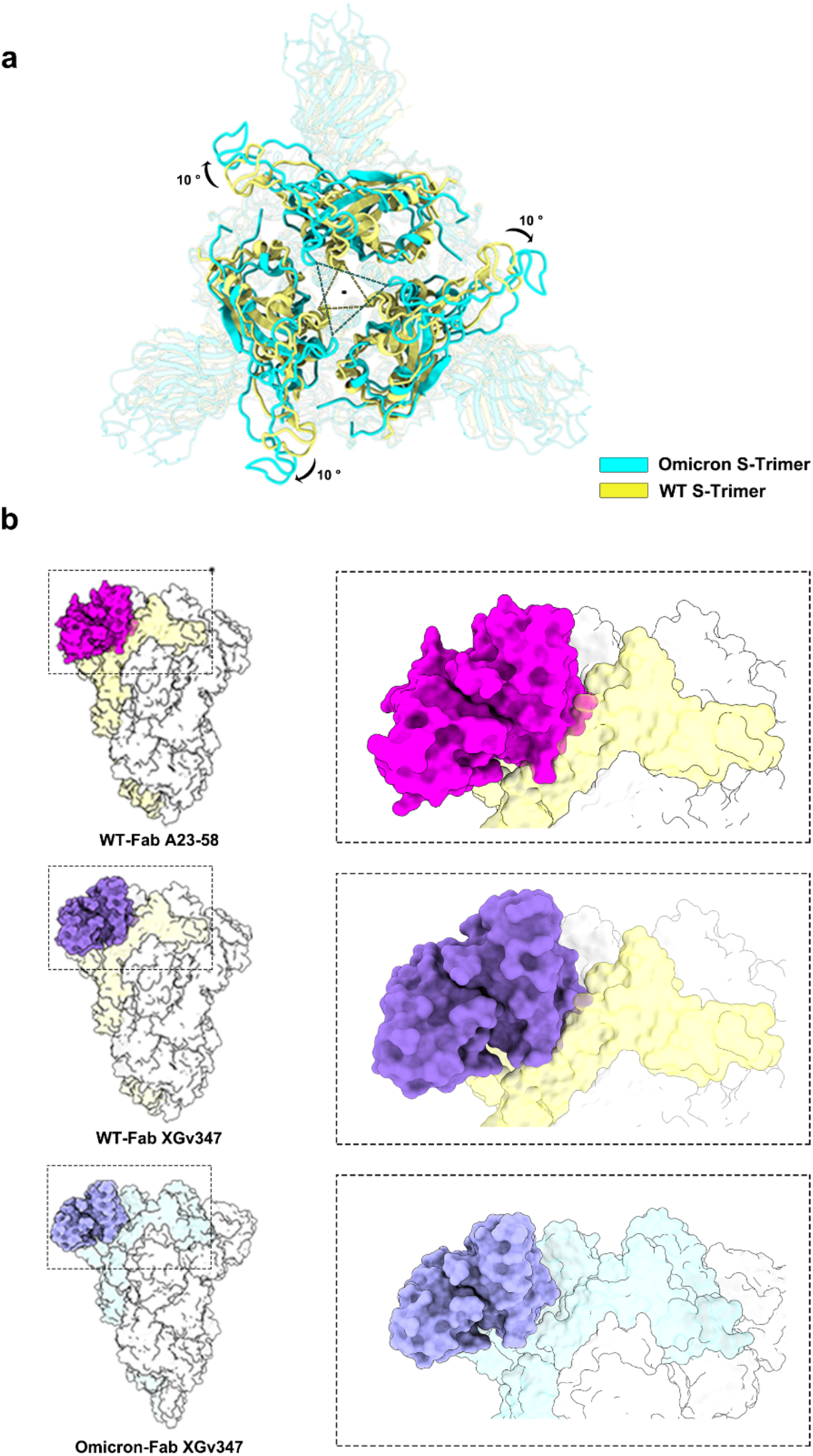
Mechanism of XGv347 binding to 3 closed RBD. **a,** Superimposition of Omicron to WT Spike Trimer. Omicron Spike is colored in cyan and wild-type Spike is colored in yellow. **b,** Superimposition of Fab A23-58 onto Omicron and WT Spike trimer and Fab XGv347 onto WT Spike trimer and are shown as surface. Fab A23-58 is colored in magenta and XGv347 is colored in purple.

**Extended Data Fig. 7.**
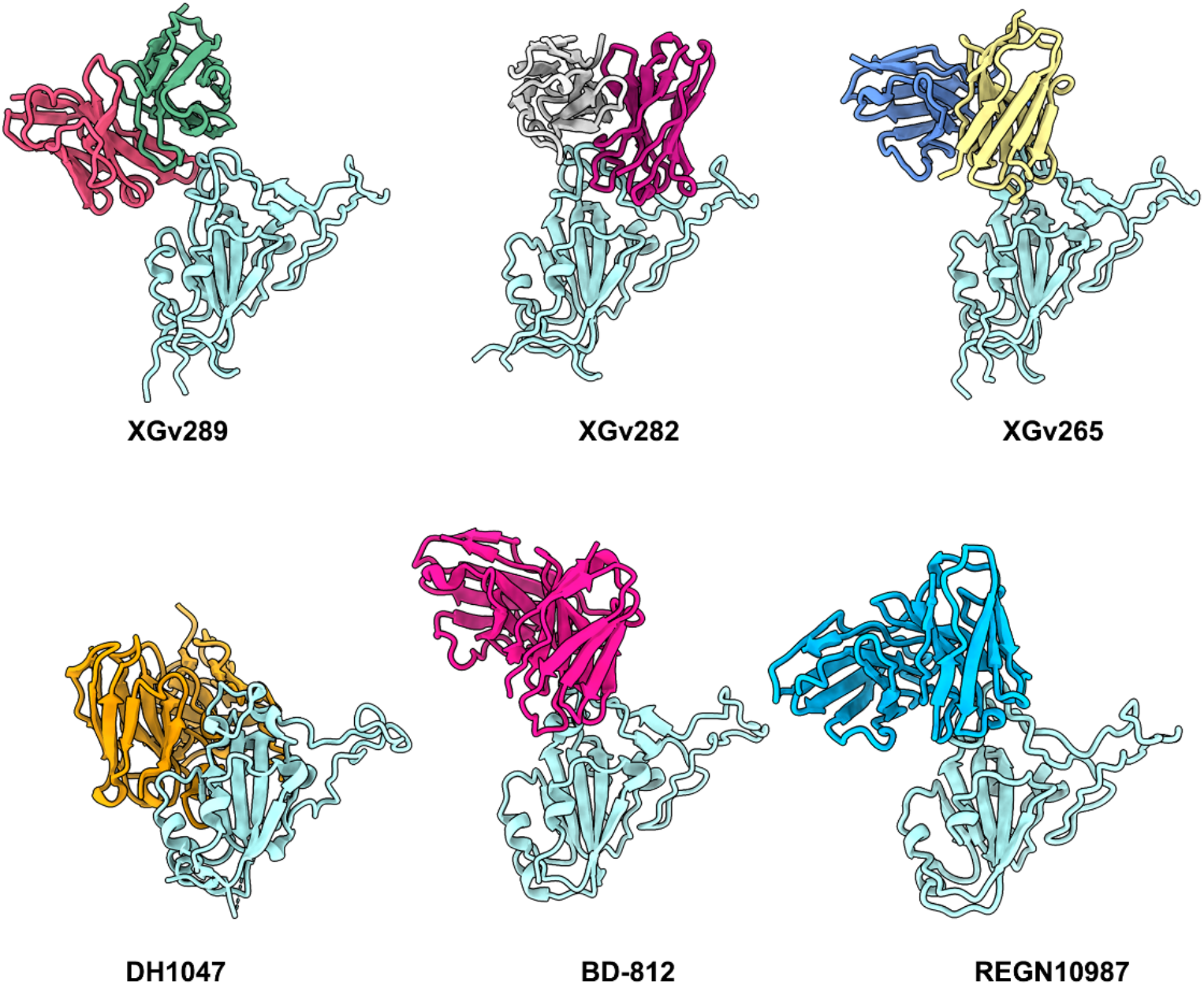
Binding modes of XGv289, 282 and 265. Binding modes of XGv289, XGv282 and XGv 265. RBD is colored in light cyan and color scheme of XGv289, XGv282 and XGv265 is the same as in Fig. 3a. DH1047, BD-812 and REGN10987 are colored in orange, deep pink and blue, respectively.

**Extended Data Fig. 8.**
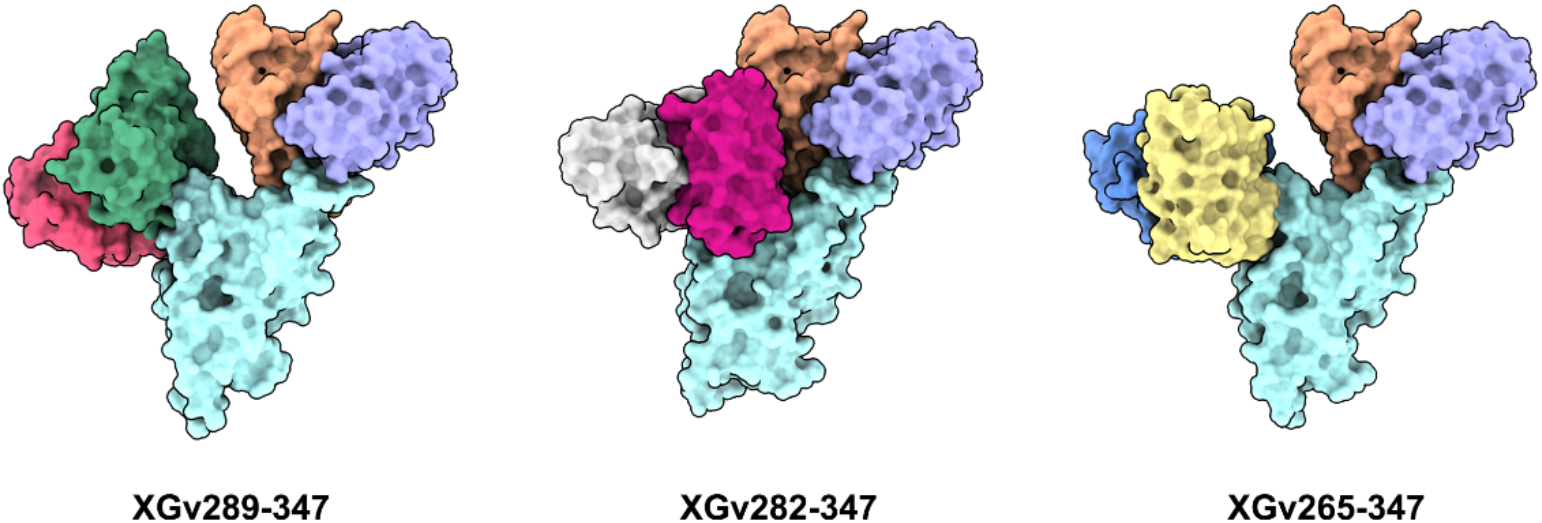
Potential cocktail representation. XGv265, XGv282 and XGv289 are superimposed onto XGv347 and all structure are shown as surface.

**Extended Data Table. 1.** Statistics for Cryo-EM data collection, refinement, and validation.

| Name | XGv347<br>in<br>complex<br>with S<br>(state 1) | XGv347<br>in<br>complex<br>with S<br>(state 2) | XGv347<br>in<br>complex<br>with S<br>(state 3) | XGv347-RBD-interface | XGv289<br>in<br>complex<br>with S | XGv289-RBD-interface | XGv282<br>in<br>complex<br>with S | XGv265<br>in<br>complex<br>with S | XGv2657-RBD-interface |
| --- | --- | --- | --- | --- | --- | --- | --- | --- | --- |
| **Data collection** |  |  |  |  |  |  |  |  |  |
| Microscope | FEI<br>Titan<br>Krios | FEI<br>Titan<br>Krios | FEI Titan<br>Krios | FEI Titan<br>Krios | FEI Titan<br>Krios | FEI Titan<br>Krios | FEI<br>Titan<br>Krios | FEI Titan<br>Krios | FEI Titan<br>Krios |
| Camera | K3 | K3 | K3 | K3 | K3 | K3 | K2 | K3 | K3 |
| Voltage (kV) | 300 | 300 | 300 | 300 | 300 | 300 | 300 | 300 | 300 |
| Total dose (e<sup>-</sup>/Å) | 60 | 60 | 60 | 60 | 60 | 60 | 60 | 60 | 60 |
| Symmetry<br>imposed | C1 | C3 | C1 | C1 | C1 | C1 | C1 | C1 | C1 |
| **Data Processing** |  |  |  |  |  |  |  |  |  |
| Micrographs<br>(Total) | 5,014 | 5,014 | 5,014 | 5,014 | 3,955 | 3,955 | 2,631 | 3,752 | 3,752 |
| Micrographs<br>(Used) | 4,561 | 4,561 | 4,561 | 4,561 | 3,864 | 3,864 | 2,598 | 3,654 | 3,654 |
| Particles selected | 232,0416 | 2320,416 | 2,320,416 | 2,320,416 | 2,332,045 | 2,332,045 | 756,508 | 1,302,103 | 1,302,103 |
| Particles<br>included in final<br>reconstruction | 269,947 | 85,822 | 105,455 | 527,413 | 401,170 | 401,170 | 119,800 | 138,359 | 138,359 |
| **Reconstruction** |  |  |  |  |  |  |  |  |  |
| Pixel size (Å) | 1.07 | 1.07 | 1.07 | 1.07 | 1.07 | 1.07 | 1.04 | 1.07 | 1.07 |
| Defocus range<br>(μm) | -1.5--2.5 | -1.5--2.5 | -1.5--2.5 | -1.5--2.5 | -1.5--2.5 | -1.5--2.5 | -1.5--<br>2.5 | -1.5--2.5 | -1.5--2.5 |
| Resolution (Å)<br>(FSC = 0.143) | 3.3 | 3.3 | 3.7 | 3.5 | 3.6 | 3.8 | 3.8 | 3.5 | 4 |
| **Model**<br>**Refinement** |  |  |  |  |  |  |  |  |  |
| Clashscore | 9.48 | 9.82 | 11.25 | 6.98 | 11.95 | 8.46 | 13.06 | 9.59 | 8.2 |
| Rotamer outliers<br>(%) | 0.00 | 0.03 | 0.00 | 0 | 0.03 | 0 | 0.00 | 0.03 | 0 |
| Molprobability score | 1.90 | 1.94 | 1.95 | 1.88 | 2.09 | 1.93 | 1.94 | 1.86 | 1.76 |
| **Ramachandran**<br>**statistics (%)** |  |  |  |  |  |  |  |  |  |
| Most favored<br>(%) | 94.06 | 93.46 | 94.31 | 91.76 | 91.69 | 92.49 | 95.42 | 94.93 | 95.51 |
| Allowed (%) | 5.83 | 6.51 | 5.67 | 8.24 | 8.23 | 7.51 | 4.56 | 5.07 | 4.49 |
| Outliers (%) | 0.11 | 0.03 | 0.03 | 0 | 0.08 | 0 | 0.03 | 0 | 0 |
| **R.m.s.deviation** |  |  |  |  |  |  |  |  |  |
| Bond lengths (Å) | 0.003 | 0.01 | 0.003 | 0.003 | 0.004 | 0.003 | 0.002 | 0.002 | 0.005 |
| Bond angles (°) | 0.714 | 0.666 | 0.567 | 0.7 | 0.62 | 0.624 | 0.557 | 0.576 | 0.72 |

**Extended Data Table. 2.**
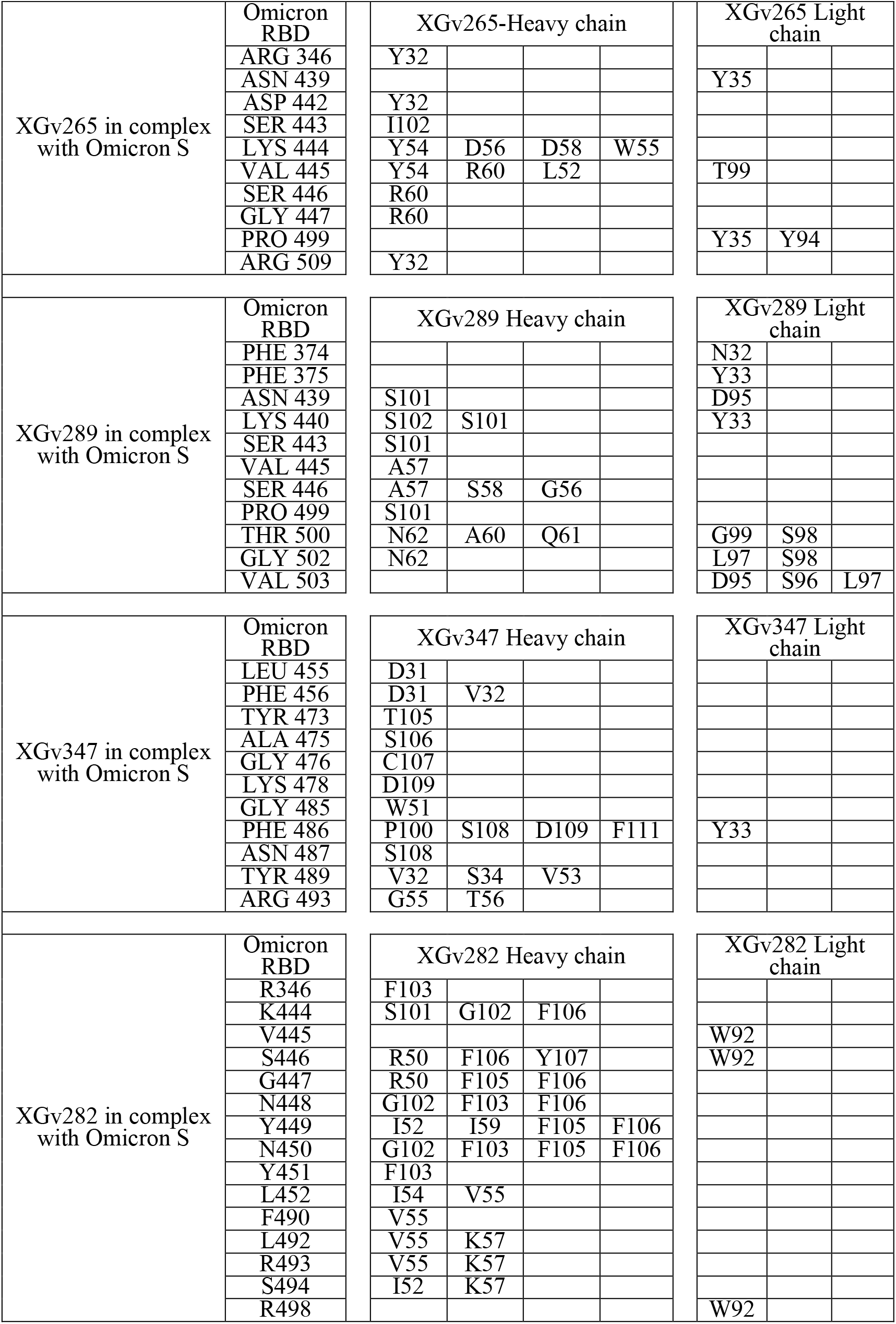
Residues of Fabs interacting with the Omicron SARS-CoV-2 S trimer at the binding interface (d < 4 Å)

**Extended Data Table. 3.** Statistics for Molecular Dynamics.

| | SARS-CoV-2 Variant | KD (nM) | $\Delta G$ (kcal/mol) | $\Delta\Delta G$ (kcal/mol) | No.(residue<sub>TOTAL</sub>) | No.(residue<sub>RB</sub>) | No.(residue<sub>fab</sub>) | No. (HB or SB) | No.(nonpolar residue<sub>RBD</sub>) | No.(nonpolar residue<sub>fab</sub>) |
| --- | --- | --- | --- | --- | --- | --- | --- | --- | --- | --- |
| XGv265 | WT | 1.475 | -3.99 | -0.96 | 21 | 10 | 11 | 14 | 4 | 7 |
|  | Omicron | 28.52 | -3.03 |  | 21 | 10 | 11 | 9 | 3 | 7 |
| XGv282 | WT | 0.8612 | -3.79 | -1.82 | 28 | 15 | 13 | 13 | 7 | 8 |
|  | Omicron | 4.096 | -1.97 |  | 19 | 8 | 11 | 7 | 3 | 8 |
| XGv289 | WT | 1.287 | -5.94 | -0.79 | 21 | 9 | 12 | 16 | 4 | 6 |
|  | Omicron | 14.17 | -5.15 |  | 26 | 11 | 15 | 12 | 3 | 4 |
| XGv347 | WT | 0.1518 | -5.42 | -0.14 | 23 | 10 | 13 | 15 | 4 | 5 |
|  | Omicron | 6.812 | -5.28 |  | 26 | 11 | 15 | 9 | 4 | 5 |

## Notes

### Competing Interest Statement

The authors have declared no competing interest.

### Summary of Updates

author affiliations and acknowledgement updated

